# Reconstructing 50 million years of *Xenopus borealis* evolution: three temporal strata of DNA rearrangements and persistent sex chromosome homomorphism

**DOI:** 10.64898/2026.03.05.709847

**Authors:** Barbora Bergelová, Nicola R. Fornaini, Tereza Tlapáková, Jiří Vávra, Marie Pleváková, Halina Černohorská, Svatava Kubíčková, Vladimír Krylov, Ben J. Evans, Martin Knytl

## Abstract

Genomic rearrangements are fundamental drivers of biodiversity, yet dynamics of structural evolution following polyploidization remain poorly understood. Genus *Xenopus* provides a valuable tool to study these phenomena. Utilizing the diploid *X. tropicalis* as a reference, we employed cytogenetic and genomic mapping to track the structural evolution of the allotetraploids *X. borealis* and *X. laevis* across a 50-million-year timeline. Based on chromosome morphometrics and C-banding patterns, we characterized the *X. borealis* pseudotetraploid karyotype (2*n* = 4*x* = 36), localizing the nucleolus organizer region (NOR) to chromosome 5L, U1 and U2 small nuclear DNAs to 1S and 8L, and 5S rDNA to nearly all chromosomes. Our analysis revealed 17 genomic rearrangements distributed within three temporal strata: ancestral (50-35 Mya), intermediate (35-15 Mya), and recent (< 15 Mya). Although we categorized chromosome 9/10 fusion as an ancestral rearrangement, the 2/9 translocation previously identified in *X. mellotropicalis* was absent in both studied allotetraploids. Furthermore, we tested for sex-specific structural polymorphism on the *X. borealis* W chromosome. Despite a large region of recombination suppression between the W and Z, no inversions were detected, indicating persistent sex chromosome homomorphism. Results are consistent with the expectation that tandem repeats such as NORs follow an asymmetric trajectory driven by a ’jumping’ mechanism and biased deletion, whereas small nuclear DNA loci are governed by copy number reduction-expansion dynamics. These findings indicate that structural rearrangements in *Xenopus* were not limited to punctuated bursts immediately following whole-genome duplication; rather, they accumulated over a prolonged evolutionary history, affecting the entire polyploid complement.

## 1. Introduction

Genomic rearrangements represent fundamental shifts in DNA architecture that reshape chromosomal landscape through the addition, deletion, movement, or reordering of genome segments compared to the ancestral position. These structural alterations serve as significant drivers of biodiversity and set the tempo of evolution, i.e., how fast those changes occur relative to a time scale (Knytl et al, submitted) [1,2]. Therefore, they have become a central subject of evolutionary research for nearly a century [3,4].

Genomic variation can be studied at multiple scales, ranging from single nucleotides to entire genomes. At the finest scale, single nucleotide polymorphisms (SNPs) are widely utilized in population genetics to assess divergence between populations [5,6] and to investigate sex-determination mechanisms via sex-specific SNPs [7,8]. At the structural level, some individuals may differ by presence or absence of a specific portion of a gene - a phenomenon known as structural sex-specific variation or allelic diversification [9–11]. Furthermore, presence or absence of entire genes can result from gene duplication often followed by movement to a new genomic location (translocation), where the gene acquires a novel sex-related function [12–14]. Another type of rearrangement is reordering of a certain chromosome segment, i.e., chromosomal inversion, which may suppress recombination between homologous chromosomes in heterozygous individuals [15]. Recombination suppression is a crucial driver of sex chromosome evolution, allowing nascent X/Y or Z/W to diverge in content [16,17]. Additionally, inversions can also contribute to speciation by accumulating differences between closely related, sympatric species, thereby creating reproductive barriers and sterility [18,19].

The most profound level on which structural/genomic variation can occur is the whole-chromosome complement, known as polyploidization or whole-genome duplication. Polyploidization is now recognized as a key driver of fish and amphibian biodiversity, with nearly one hundred independent polyploidization events documented, including that ancestral for all vertebrates (Knytl et al, submitted). Although historically thought to be redundant and an evolutionary dead-end, polyploidy has the potential to provide adaptive potential, facilitating geographical invasion and survival in changing environmental conditions [20–22]. However, the role of polyploidization in sex determination remains poorly understood, warranting further research attention.

### 1.1. Pipid frogs as a model system for genomic variation

Given at least 11 independent polyploidization events evidenced within the amphibian family Pipidae [23–26], this group represents a valuable model for studying evolution by polyploidization. *Xenopus* is a genus with an exceptionally high number of polyploids, in which all but one species are polyploid [27]. Ploidy levels range from diploid to dodecaploid, spanning from 2*n* = 2*x* = 20 to 2*n* = 12*x* = 108 chromosomes [27]; here, *n* denotes the gametic (haploid) chromosome number of the extant species, and *x* refers to the haploid number of chromosomes of the most recent diploid ancestor of the extant species. All *Xenopus* polyploids behave like cytological diploids, forming meiotic bivalents rather than multivalents and gametes with reduced genetic content due to the rediploidization of the polyploid genome [27]. The formation of meiotic bivalents is an indication of allopolyploid origin (hybridization between/among distinct diploid ancestors) rather than autopolyploidy (multiplication within a single ancestor) [28]. Also, allopolyploids have their genome fragmented into units inherited from lower ploidy ancestors, known as subgenomes [29–33].

The diploid species *Xenopus tropicalis* (2*n* = 2*x* = 20) belongs to subgenus *Silurana* along with other three allotetraploid species with 2*n* = 4*x* = 40, *X. calcaratus*, *X. mellotropicalis*, and *X. epitropicalis* [24]. As the only diploid species, *Xenopus tropicalis* serves as a reference material and potential parental ancestor for evolutionary and phylogenetic enigmas [34,35]. At least two allotetraploidization events happened in subgenus *Silurana*, the one giving rise to *X. calcaratus*, and the other forming a common clade of *X. mellotropicalis* and *X. epitropicalis*. These events were estimated to occur recently, less than 10 mya [36]. Genomes of *Silurana* polyploids are consisting of a- and b-subgenomes, which are distinguishable based on cytogenetic methods and Sanger sequencing [23,34,35] even despite their recent evolutionary split of ∼10 Mya [24,36].

Within subgenus *Xenopus* (which diverged from *Silurana* ∼50 Mya; [30]), species exhibit tetraploidy (2*n* = 4*x* = 36), octoploidy (2*n* = 8*x* = 72), and dodecaploidy (2*n* = 12*x* = 108) [27]. In these species, subgenomes are designated L and S and exhibit asymmetric genomic evolution. Specifically, the L-subgenome displays higher syntenic conservation and fewer chromosomal rearrangements than the S-subgenome when compared to the reference genome structure of *X. tropicalis* [23,30]. The tetraploid species *X. laevis* (2*n* = 4*x* = 36) serve as a cornerstone vertebrate model for developmental and evolutionary research [33,37–40]. *Xenopus borealis* serves as an emerging model for evolutionary biology and sex chromosomes [41,42]. Together with the diploid *X. tropicalis*, they represent a useful tool for comparative studies centered on genome evolution by polyploidization and structural variation [14,16,30]. Furthermore, chromosome-scale genome assemblies for all these three species are available [30,41,43].

Following polyploidization, duplicated gene copies often follow divergent evolutionary trajectories [30,42], and thus the process may be accompanied by subgenome-specific chromosomal rearrangements as a consequence of the rediploidization process [36]. A key genomic event is the fusion of ancestral chromosomes 9 and 10, which is hypothesized to have occurred in the common ancestor of subgenus *Xenopus* [30]. This hypothesis is supported by conserved number of chromosomal echelons across the subgenus, as all species of this group share 36-chromosomal karyotypes or multiples thereof [27]. High-resolution whole-genome sequencing and cytogenetic gene mapping have further refined the fusion interpretation by characterizing breakpoints and fusion junctions [30,44]. Beyond this shared fusion, a lineage-specific rearrangement has been detected, specifically the translocation of a heterochromatic block from ancestral chromosome 9b to 2b in *X. mellotropicalis* [34,35]. It is unknown whether the translocation happened in a diploid or polyploid ancestor.

Chromosomal inversions can be identified by synteny mapping and collinearity analysis [40,43,45,46]. In *Xenopus laevis*, at least five such inversions occurred on chromosomes 2S, 3S, 4S, 5S and 8S [30,40]. Notably, all these inversions are situated in the S-subgenome, underscoring an increased susceptibility of the S-subgenome to structural reorganization compared to the stable L-subgenome [30]. It remains to be determined whether additional unidentified inversions in *Xenopus* contribute to recombination suppression and the subsequent divergent evolution of sex chromosomes typically favored by natural selection.

### 1.2. Cytogenetics as a tool for chromosome identification

Measurement of chromosome arms is an effective technique with which to identify homologous chromosome pairs and arrange them into karyotypes [47–50]. In all *Silurana* species, each chromosome has been clearly distinguishable based on mensural analysis [34,35,51]. However, some non-homologous chromosomes in subgenus *Xenopus* are very similar in terms of length and arm ratio, and therefore additional cytogenetic techniques may be used to identify nearly identical chromosomes between each other (see e.g., [44,47]). Chromosome banding (C-, G-, Cy3-) has been introduced for *Xenopus* chromosome identification. However, C- and Cy3- banding detected blocks of constitutive heterochromatin on one chromosome pair, at the same regions where nucleolar organizer regions (NORs) are formed (reviewed in [27]). G-banding did not show any specific signals (unpublished data). Another method that can be used for identifying nearly identical *Xenopus* chromosomes is fluorescence *in situ* hybridization (FISH) utilizing probes targeting tandem arrays, such as U1 and U2 small nuclear DNA (snDNA) and 5S and 28S ribosomal DNA (rDNA) [36,44,52]. None of these four methods have been used in *X. borealis*, except for 5S rDNA FISH showing signals on the long arms of nearly all chromosomes in the karyotype [52].

### 1.3. Goals

In this study, we provide a comprehensive characterization of the karyotype of the allotetraploid species *Xenopus borealis* and investigate subgenome-specific rearrangements—including inversions, fusions, and translocations—within the L- and S-subgenomes of *X. borealis* and *X. laevis*. By integrating chromosome morphometrics, C-banding, and a suite of FISH techniques with whole-genome synteny mapping, we delineate the structural evolution of these tetraploid genomes. Furthermore, we explore the structural divergence of the *X. borealis* sex chromosomes by assessing whether a putative inversion within the sex-determining locus has facilitated their differentiation.

## 2. Material and methods

### 2.1. Primary cell cultures and metaphase spread preparations

*Xenopus borealis* animals were originally obtained from the Institute of Zoology at the University of Geneva (Switzerland). All individuals were bred at Charles University, Faculty of Science, Prague, Czech Republic. Briefly, tadpoles were anesthetized, hind limbs removed and homogenized [53] in a cultivation medium [34] and modified according to Knytl et al. [35]. The explants were then cultivated at 29.5°C with 5.5% CO_2_ for five days without disturbance. The medium was then changed every day for one week. The first and next passages were performed with trypsin-ethylenediaminetetraacetic acid [34]. After two weeks, Antibiotic-Antimycotic (100X) solution (Gibco™ by Thermo Fisher Scientific, Waltham, MA, USA) was removed from the cultivation medium.

Chromosomal suspensions were prepared from two males and two females according to Khokha et al. [54] with minor changes [55] and then stored in fixative solution (methanol: acetic acid, 3:1, v/v) at -20°C. For laser microdissection, a fresh metaphase suspension was dropped onto a polyethylene-naphthalene membrane. For cytogenetic analysis, a chromosome suspension was dropped onto a slide [56]. Chromosome preparations were aged at -20°C for at least one week with the exception of that for FISH with tyramide signal amplification (FISH-TSA), in which the chromosome suspension was dropped and directly followed by further experimental procedures. For each experiment, mitotic metaphase spreads were counterstained with ProLong™ Diamond Antifade Mountant with the fluorescent 4’,6-diamidino-2-phenylindole, DAPI stain (Invitrogen by Thermo Fisher Scientific). From ten to twenty metaphase spreads were analyzed per each probe. Microscopy and processing of metaphase images was conducted using Leica Microsystem (Wetzlar, Germany) [44].

### 2.2. Laser microdissection and whole chromosome painting

We used whole chromosome painting (WCP) probes generated by laser microdissection of *X. tropicalis* chromosomes from the study Knytl et al. [35]. Probes were labeled using GenomePlex WGA Reamplification Kit, WGA3 (Sigma-Aldrich, St. Louis, MO, USA) either with digoxigenin-11-dUTP [55] or biotin-16-dUTP [35] (both Jena Bioscience, Jena, Germany). Control painting FISH on *X. tropicalis* and cross-species painting FISH (Zoo-FISH) on *X. borealis* chromosomes were carried out as described by Krylov et al. [55] with minor modifications as detailed in Knytl et al. [34]. Autoclaved *X. tropicalis* genomic DNA was used as a competitor (blocking DNA) according to Bi and Bogart [57]. Fluorescent signal of digoxigenin and biotin labeled probes was visualized by anti-digoxigenin-rhodamine (Roche, Basel, Switzerland) and streptavidin, Cy3 (Invitrogen, Camarillo, CA, USA), diluted with blocking reagents as used in double-colour FISH with ribosomal probes in Knytl et al. [34].

### 2.3. C-banding, and FISH with snDNA and rDNA probes

The C-banding protocol followed Rábová et al. [58], with the modifications described in Knytl et al. [34]. In order to generate probes for repetitive FISH (the U1 and U2 snDNA probes; 5S and 28S rDNA probes), *X. tropicalis* was used as a template for amplification of U1 and U2 snDNA, and 5S rDNA regions. *Xenopus laevis* was used as a template for 28S rDNA probe. Total gDNA was extracted from the web tissue of adult frogs using the DNeasy Blood & Tissue Kit (Qiagen, Hilden, Germany) according to manufacturer’s instructions. Conditions for PCR amplification were used according to PPP Master Mix (Top-Bio, Prague, Czech Republic) supplier recommendations. Primers used for U1 and U2 snDNA amplification are listed in Table 1 of Fornaini et al. [36]. Primers used for 5S and 28S rDNA mapping are shown in Table S1 of Knytl et al. [35]. PCR conditions for the U1 and U2 snDNA amplifications were taken from Fornaini et al. [36]. The 5S and 28S rDNA loci were amplified as detailed in Knytl and Fornaini [59]. PCR labeling of all probes (U1, U2, 5S, and 28S) with digoxigenin-11-dUTP and biotin-16-dUTP (both Jena Bioscience) is detailed in Knytl and Fornaini [59]. Digoxigenin-11-dUTP was used for U2 and 5S labeling. Biotin-16-dUTP was used for U1 and 28S labeling. *Xenopus tropicalis* U1, U2, and 5S probes were hybridized with chromosome spreads of *X. borealis*. The contents of the hybridization mixture, denaturation, and the subsequent overnight hybridization were conducted as it described in rDNA FISH protocol [35]. Post-hybridization washing and blocking reactions were performed as described for painting FISH in Krylov et al. [55]. Probe visualization was performed as protocol [34]. For sequential FISH mapping, slides were destained according to Fornaini et al. [36] and subsequently used for additional round of FISH experiment.

**Table 1.**
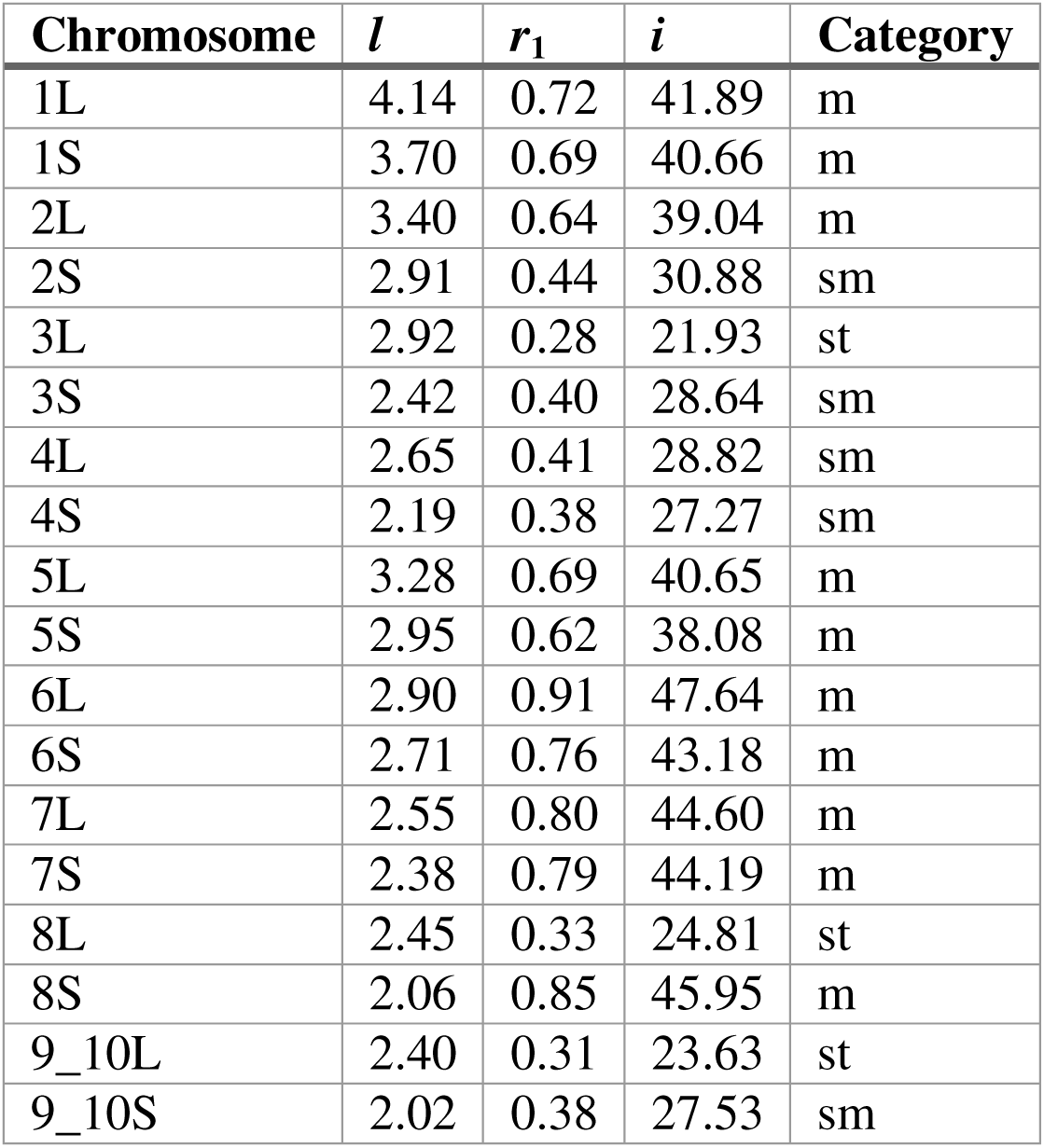
*Xenopus borealis* chromosome characteristics for each haploid chromosome. shown in the medians of chromosome length (*l*), p/q arm ratio (*r*_1_), and centromeric index (*i*).

| Chromosome | $l$ | $r_1$ | $i$ | Category |
| --- | --- | --- | --- | --- |
| 1L | 4.14 | 0.72 | 41.89 | m |
| 1S | 3.70 | 0.69 | 40.66 | m |
| 2L | 3.40 | 0.64 | 39.04 | m |
| 2S | 2.91 | 0.44 | 30.88 | sm |
| 3L | 2.92 | 0.28 | 21.93 | st |
| 3S | 2.42 | 0.40 | 28.64 | sm |
| 4L | 2.65 | 0.41 | 28.82 | sm |
| 4S | 2.19 | 0.38 | 27.27 | sm |
| 5L | 3.28 | 0.69 | 40.65 | m |
| 5S | 2.95 | 0.62 | 38.08 | m |
| 6L | 2.90 | 0.91 | 47.64 | m |
| 6S | 2.71 | 0.76 | 43.18 | m |
| 7L | 2.55 | 0.80 | 44.60 | m |
| 7S | 2.38 | 0.79 | 44.19 | m |
| 8L | 2.45 | 0.33 | 24.81 | st |
| 8S | 2.06 | 0.85 | 45.95 | m |
| 9_10L | 2.40 | 0.31 | 23.63 | st |
| 9_10S | 2.02 | 0.38 | 27.53 | sm |

### 2.4. Single-copy gene mapping using FISH-TSA

RNA from *X. borealis* brain, testes, and spleen was extracted using E.Z.N.A. Total RNA Kit I (Omega Bio-tek; Norcross, GA, USA). cDNA was prepared from 200 ng of RNA by means of first strand reverse transcription SuperScript™ III Reverse Transcriptase (Invitrogen by Thermo Fisher Scientific). A protein coding cDNA was amplified using PPP Master Mix (Top-Bio, Prague, Czech Republic) according to manufacturer’s instructions (annealing temperature 54-59°C). Eight genes were amplified:

*choline/ethanolamine phosphotransferase1* (*cept1*), *glycogenin 2* (*gyg2*), *fibronectin 1* (*fn1*), *NADH: ubiquinone oxidoreductase core subunit S1* (*ndufs1*), *splicing factor 3b subunit 1* (*sf3b1*), *nuclear receptor subfamily 5 group A member 1* (*nr5a1* aka *sf-1*), *androgen receptor* (*ar*), and *SRY-box 3* (*sox3*). Primer sequences are listed in Supplementary material, Table S1. Post PCR purification was conducted as detailed in Knytl and Fornaini [59]. The cDNA amplicons were cloned using the TOPO-TA cloning kit and One Shot™ TOP10 Chemically Competent *E. coli* (both Invitrogen by Thermo Fisher Scientific). Plasmid DNA was isolated from bacterial colonies was isolated from 4–5 bacterial colonies for each gene using the E.Z.N.A. Plasmid DNA Mini Kit I (Omega Bio-tek), checked by electrophoresis, and sent for sequencing. Sanger sequences were compared with sequences from *X. borealis* using Geneious Prime, version 2025.2.2. The cDNA sequences were deposited in the GenBank database (Supplementary material, Table S1). Subsequently, the loci were re-amplified and purified [59]. The probes were labeled with digoxigenin-11-dUTP (Jena Bioscience) using DecaLabel DNA Labeling Kit (Thermo Scientific by Thermo Fisher Scientific). The FISH-TSA protocol was adopted from Krylov et al. [60] with minor modifications described in Knytl et al. [61]. FISH probes were labeled with digoxigenin-11-dUTP (Roche, Mannheim, Germany), signal was detected by antidigoxigenin-POD, Fab fragments (Roche), and amplified using the TSA TM-Plus Tetramethylrhodamine System Kit (NEL742001KT, PerkinElmer, Inc., Waltham, MA, USA). Some metaphase spreads were used for sequential FISH-TSA mapping followed by U2 snDNA FISH.

### 2.5. Measurement and identification of *X. borealis* chromosomes

A total of 19 *X. borealis* metaphase figures (8 male, 11 female metaphases) were analyzed from 4 individuals (2 males, 2 females). Half of the metaphases selected for the identification of individual chromosomes was stained with 5% Giemsa/PBS solution (v/v). The rest were stained with DAPI during FISH experiments. Arms were measured in pixels using Adaptive Contrast Control Scientific Image Analyzer software (Sofo ACC 6.2, Brno, Czech Republic). The p and q arm lengths were quantified as described in Knytl and Fornaini [59]. To identify each chromosome, we analyzed chromosomal length (*l*), arm ratio (*r*_1_) [62], centromeric index (*i*), and p/q arm ratio (*r*_2_) [63] according to formulas shown in Knytl et al. [35]. Chromosomal nomenclature was taken from Matsuda et al. [47]. Each individual chromosome was also assigned a chromosomal category based on the *i* value. If the *i* value was equal to or greater than 37.5, the chromosome was categorized as metacentric. If the *i* value was equal to or higher than 25 and lower than 37.5, the chromosome was categorized as submetacentric, and if the *i* value was equal to or greater than 12.5 and lower than 25, the chromosome was categorized as subtelocentric. Data were analyzed in R software for statistical computing, V 4.5.2 (R Core Team 2025) using the ggplot2 and ggpubr packages. Two-way analysis of variance (two-way ANOVA) was performed to test whether the *l* and *i* values differ significantly between (1) L and S homoeologous chromosomes regardless of sex; (2) males and females, and (3) whether there is an interaction effect between chromosome type and sex. Whether there is an interaction effect, Tukey’s test found all interaction possibilities and the most important was the interaction between males and females within the same chromosome (e.g., Chr8L males vs Chr8L females). Original R scripts were modified from Knytl and Fornaini [59]. The complete computational workflow, including scripts for morphometric calculations, table and plot generation, as well as original metaphase images, are publicly available on https://github.com/martinknytl/2025_borealis_karyotype.

### 2.6. Whole-genome synteny mapping within *X. tropicalis*, *X. borealis*, and *X. laevis*

Annotated coding sequences (CDS) for *X. tropicalis* v10.0 [43] and *X. laevis* v10.1 [30] were downloaded from the database. CDS fragments shorter than 200 bp were filtered out based on GFF3 annotations, and FASTA sequences for the retained CDS were generated using BEDTools. Separate L- and S-subgenomes were extracted from the *X. borealis* v1 [41] and *X. laevis* v10.1 [30] genome assemblies using BEDTools, and local databases were constructed using makeblastdb. The annotated *X. tropicalis* CDS were aligned to the *X. borealis* and *X. laevis* L- and S-subgenomes using the BLASTn. Then the annotated *X. laevis* CDS were mapped to *X. borealis* L- and S- subgenomes. For synteny analysis, results were filtered to retain only the single best hit for each query based on the highest bit score. The resulting datasets were then parsed to extract the input parameters required for generating synteny plots: species and chromosome IDs, start and end positions, orientation, and total chromosome lengths for both reference and query). The complete data analysis pipeline is available on https://github.com/martinknytl/Xen_rearrangements.md/blob/main/Whole-genome_synteny. Synteny plots were generated in R software V 4.5.2 for statistical computing (R Core Team 2025) using R packages syntenyPlotteR, ggplot2, ggforce, and the function draw.microsynteny for syntenyPlotteR [46]. The circular plot was created in Synteny Portal using interactive SynCircos with a minimum size of a reference block (bp) 500,000. *Xenopus tropicalis* was selected as the reference species, *X. laevis* was used as the target species.

## 3. Results

### 3.1. C-banded karyotype

C-banding generally shows constitutive heterochromatic blocks which are associated with the rich GC content. Therefore we performed this method with the aim to find chromosome-specific GC blocks of constitutive heterochromatin, and thus to arrange *X. borealis* chromosomes into karyotype. GC-rich blocks (i.e., DAPI positive/intensive) were found on the pericentromeric region of the p arms of 1L, 1S, 2L, 5L, and 5S chromosomes (Fig. 1). Some metaphases show DAPI positive signals on 7L, and 7S as well (not shown). However, based on the GC-rich blocks we are not certain in identifying homologous and homoeologous pairs of chromosomes as the C-banding pattern identifies pericentromeric area of the p arm. Secondary constriction was detected on the p arm telomere of chromosome 5L by a DAPI negative band (Fig. 1), indicating nucleolar position. Chromosomes 8L which are the sex chromosomes in *X. borealis* [64] was not detectable by the C-banding method.

**Figure 1.**
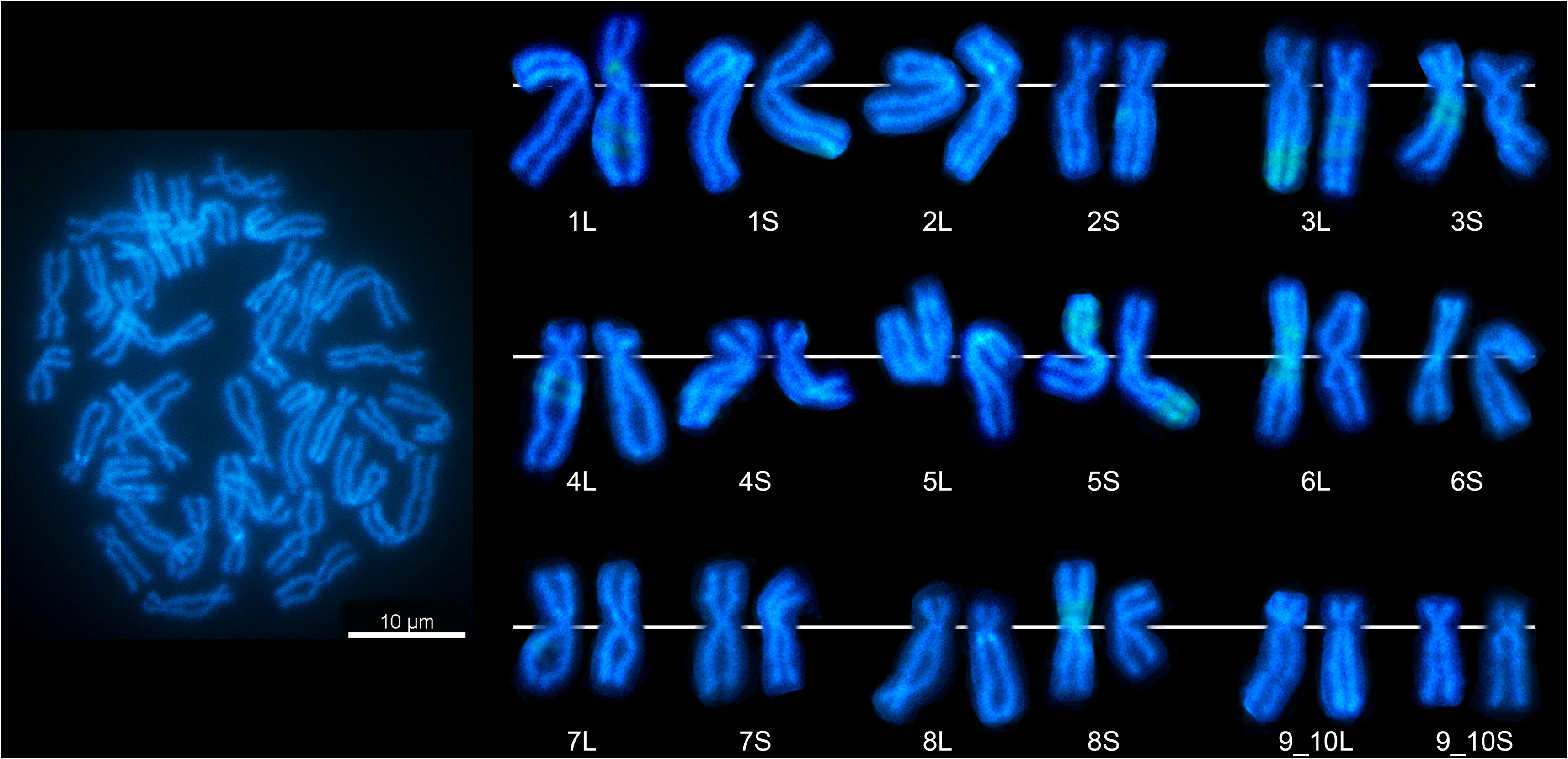
Karyotype of *X. borealis* based on the C-banding method. shows 36 chromosomes sorted into homologous pairs and nine homoeologous groups. A white line indicates the centromere position for each chromosome.

### 3.2. Zoo-FISH

Having confirmed the specificity of the *X. tropicalis* WCP probes [35], we went on to perform Zoo-FISH experiments on *X. borealis* metaphase spreads using each *X. tropicalis* WCP probes. The WCP probes derived from XTR 1, 6, 7, and 8 hybridized to whole chromosomal quartets (the homoeologous chromosome groups) without evidence of large-scale translocations as follows: WCP of XTR Chr1 to *X. borealis* chromosome 1L (XBO Chr1L) + 1S (Fig. 2a); XTR Chr6 to XBO Chr6L + 6S (Fig. 2b); XTR Chr7 to XBO Chr7L + 7S (Fig. 2c); XTR Chr8 to XBO Chr8L + 8S (Fig. 3b). In addition, WCP 6 detected a non-fluorescent interstitial part on the q arm of the XBO Chr6L (Fig. 2b, indicated by arrows in the bottom-right frame; the less condensed chromosomes shown in the frame were dissected from a different metaphase spread). This gap likely reflects a high density of repetitive sequences that were effectively masked by the competitor DNA, preventing probe hybridization in this region. The WCP 9 probe hybridized to a chromosomal segment of the q arms of XBO Chr9_10L + 9_10S (Fig. 2d), whereas the WCP 10 probe mapped to the p arms of XBO Chr9_10L + 9_10S including the pericentromeric region (Fig. 2e). Hybridization of the WCP 9 and 10 to appropriate chromosomal segments confirmed a fusion of ancestral chromosomes 9 and 10 and disappearing the centromere of ancestral chromosome 9 as the Chr9 centromere is not highlighted with WCP 9 (Fig. 2d). The remaining *X. tropicalis* probes (WCPs 2, 3, 4, and 5) did not work on *X. borealis* chromosomes for unknown reasons, but possibly due to rearrangements or other factors.

**Figure 2.**
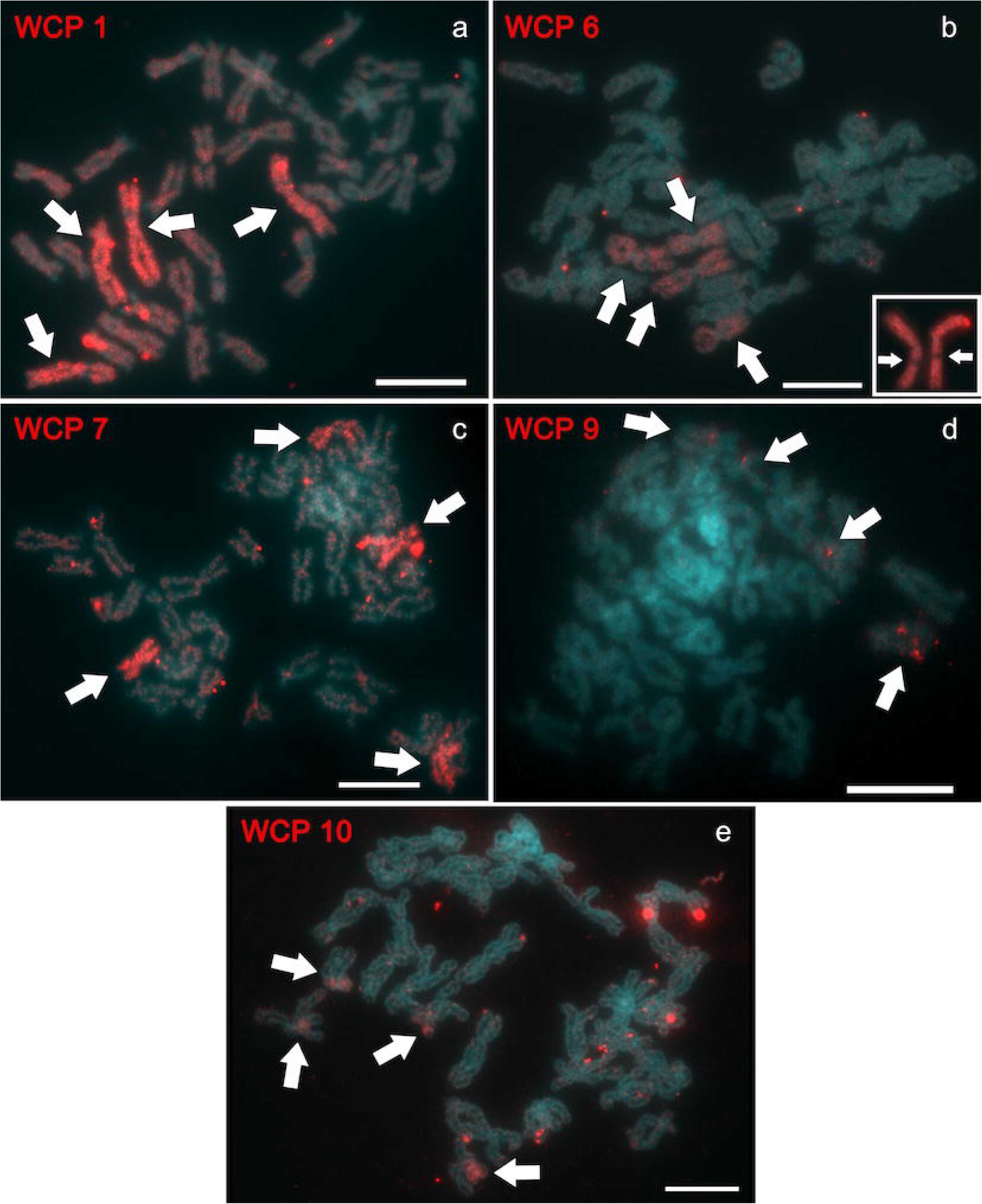
Cross-species fluorescence *in situ* hybridization with *X. tropicalis* whole-chromosome painting (WCP) probes hybridized on 36 *X. borealis* chromosomes (XBO). The WCP probes stain an appropriate group of homoeologous chromosomes. (a) WCP isolated from chromosome 1 (WCP 1) - XBO Chr1L and 1S, (b) WCP 6 - XBO Chr6L and 6S, (c) WCP 7 - XBO Chr7L and 7S, WCP 8 (shown on Figure 3) - XBO Chr8L and 8S, (d) WCP 9 – long arms of XBO Chr9_10L and 9_10S, (e) WCP 10 - short arms and pericentromeric region of XBO Chr9_10L and 9_10S.

**Figure 3.**
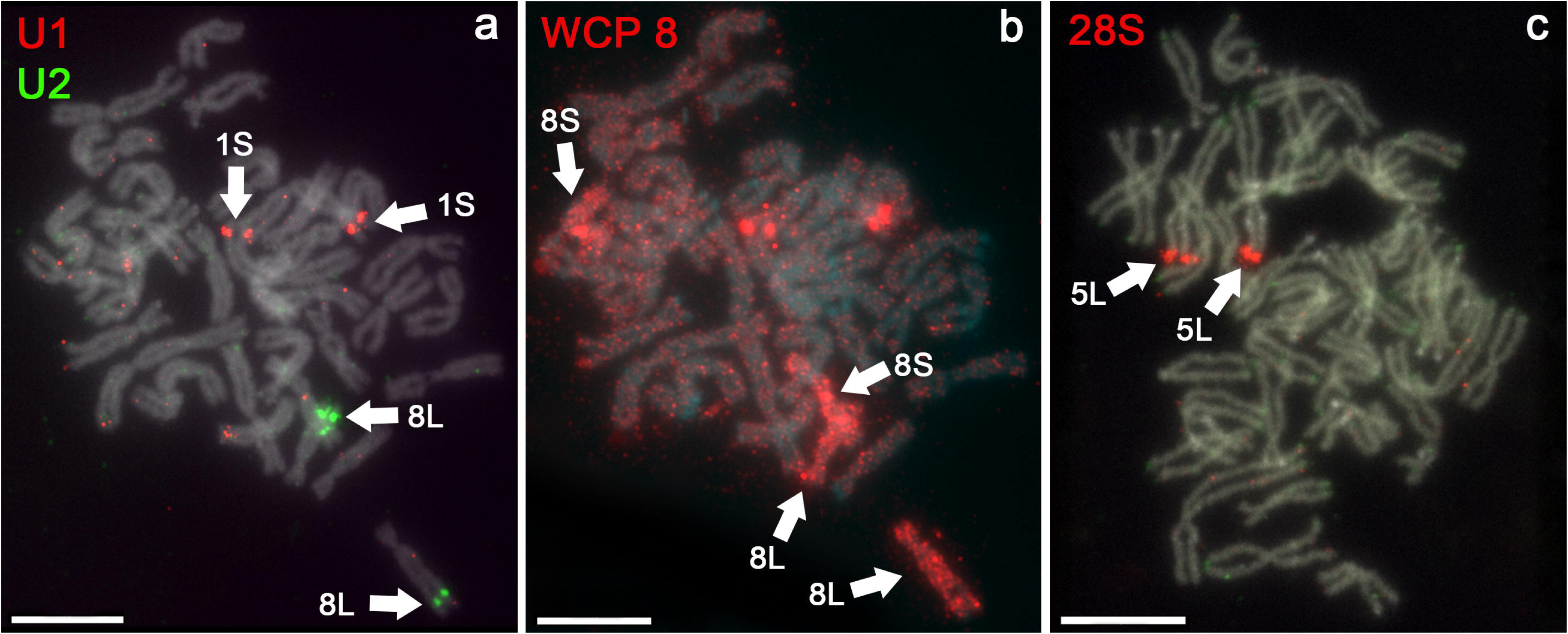
Sequential fluorescent chromosome mapping using U1 and U2 snDNA FISH, cross-species whole chromosome painting Zoo-FISH, and 5S and 28S rDNA FISH on metaphase spread of *X. borealis*. DAPI (blue-green) counter-stained metaphase spreads show all 36 chromosomes. (a) FISH with U1 (red) and U2 (green) snDNA probe identifies XBO Chr1S and XBO Chr8L, respectively. (b) FISH with the *X. tropicalis* WCP probes painted XBO Chr8L and 8S on the same metaphase plate. Red spots on chromosome 1S (unmarked) are the remaining signals from the U1 snDNA probe after destaining. (c) FISH with 28S (red) ribosomal probes identifies the p arm of XBO Chr5L where is the nucleolar secondary constriction. Scale bars represent 10 μm.

### 3.3. snDNA FISH and rDNA FISH

The U1 and U2 snRNA probes (snDNA FISH) mapped to chromosome 1S and 8L (Fig. 3a; U1 in red, U2 in green), respectively. In *X. borealis*, chromosome 8L represents the sex chromosome, characterized by a large sex non-recombining locus involving the p arm and pericentromeric region [64]. The U1 snDNA segment is located on a distal part of the q arm of chromosome 1S, while the U2 snDNA locus is situated in a distal part of the q arm of chromosome 8L, within a region that appears to be a pseudoautosomal segment. Direct confirmation that the U2 locus resides on chromosome 8L was obtained through Zooc:FISH: the *X. tropicalis* wholec:chromosome painting probe for chromosome 8 hybridized to *X. borealis* chromosomes 8L and 8S on the same metaphase spreads that had been previously stained for U2 snDNA. Co-localization of U2 snDNA with WCP XTR 8 is shown on Figure 3a, b. Beside the U2 signals on chromosome 8L some metaphases contain an additional signal on a chromosome of the same or very chromosome 4S, which has a similar morphology as 8L. It is clear that the chromosome carrying an additional U2 signal is not chromosome 8S.

Blast analysis of the U1 sequence used as a probe mapped to chromosome 1L (∼213,100,000 bp of the *X. borealis* genome), 1S, and 2S (multiple repeats of 90-97% identity), while the U2 snDNA sequences matched chromosome 8L and 8S with coordinates ∼123,800,000 and ∼105,300,000 bp of the *X. borealis* genome (repeats with 95-100% similarity, respectively). The BLAST results U2 sequence are incongruent with our cytogenetic observations, suggesting either significant inter-individual variation in U2 snDNA accumulation between the individuals used for genome sequencing and those used for cytogenetics, or a reduced copy number of those loci in the S-subgenome that falls below the detection threshold of FISH.

The 5S probe was mapped to telomeres of nearly all chromosomes. The 5S signal is very weak and better visibility of this signal is shown in a separate green channel (in Black & White mode) in Supplementary material, Figure S1. The 28S rDNA probe associated with NOR mapped to telomeres of the p arm of chromosome 5L (Fig. 3c, red). Blast analysis of the 5S and 28S rDNA sequences mapped to all chromosomes and chromosome 4L (23,315,662-23,315,875 bp of the *X. borealis* genome with 97% identity), respectively. *Xenopus borealis* is an evolutionarily tetraploid species with cytologically diploidized genome (sometimes referred as pseudotetraploid), and thus the NOR on the chromosome 5S is supposed to be lost as 28S rDNA signal was not cytogenetically localized on another chromosome than chromosome 5L. Genome mapping shows no match of the 28S locus on any chromosome in the S-subgenome. Cytogenetic analysis does not coincide with blast analysis because chromosomes 4L and 5L are non-homologous with substantially different morphology (see Discussion).

### 3.4. Testing of three large-scale rearrangements by genomic and cytogenetic mapping

We set out to assess whether: (i) The inversion affecting sex locus on chromosome 8L is present on the female W chromosome but not on the male/female Z chromosome. (ii) The fusion between ancestral chromosomes 9 and 10 occurs in the *X. borealis* lineage by the same mechanism as the fusion that occurred in *X. laevis* ancestor (sensu [30]). (iii) A translocation between chromosomes 9 and 2 is present in *X. borealis* as it is in *X. mellotropicalis*. This translocation in *X. mellotropicalis* was revealed by Zoo-FISH [34] and FISH-TSA mapping of five single-copy genes that flank the translocated region (t(9;2) translocation-associated genes) [61].

To explore (i), we hypothesize that the inversion affected the pericentromeric region of the W chromosome and exhibits sex-specific structure. Three genes were chosen for this hypothesis: *sox3.L*, *ar.L*, and *sf-1.L*. The probe cDNA sequences were mapped to the *X. borealis* genome assembly using BLASTn and the order of genes were: *sf-1.L* 12,034,730-12,036,203 bp; *ar.L* ∼18,200,000 bp; *sox3.L* 37,849,021-37,850,160 bp of the *X. borealis* genome. The genome of *X. borealis* were sequenced from a male individual and therefore the homogametic ZZ sex represents the assembly [41]. Because both male and female chromosome suspensions were available, the FISH-TSA was used for exploring the inversion in females. The FISH results did not confirm the inversion as the order of *sox3.L*, *ar.L*, and *sf-1.L* genes was identical on male and female chromosome 8L (Fig. 4; Supplementary material, Figs S2-S4), and also the order of the studied genes revealed by FISH-TSA corresponds with the order of those genes in *X. borealis* genome database. The specificity of the *sf-1* signal on the female chromosome 8L was confirmed by sequential U2 snDNA hybridization after slide destaining (Supplementary material, Fig S4c; *sf-1* in red, U2 in green).

**Figure 4.**
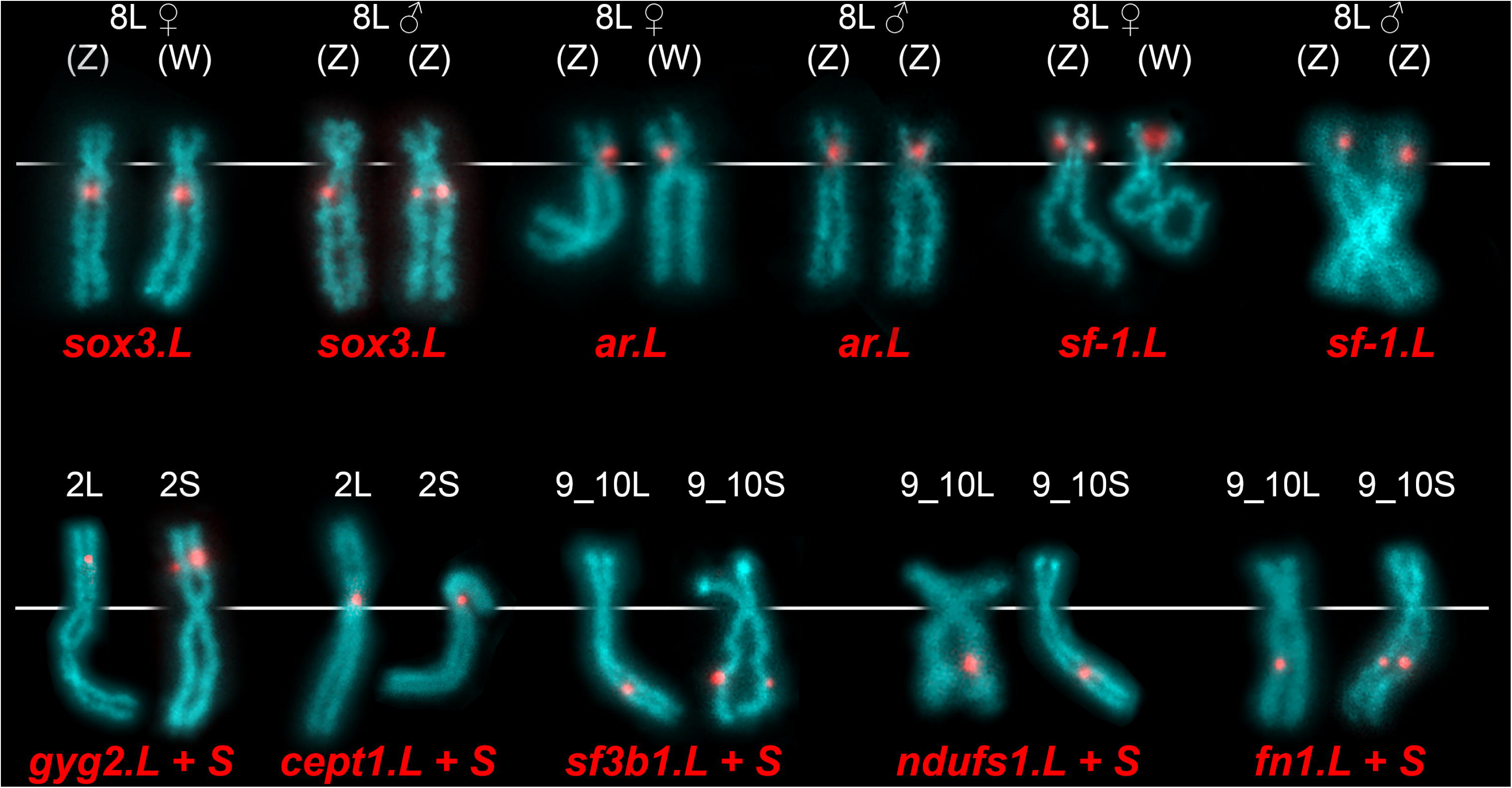
FISH coupled with tyramide signal amplification (FISH-TSA) with positive red signals on *X. borealis* chromosomes. The *gyg2.L* and *S*, *cept1* loci were localized on the p arm of XBO Chr2L and 2S, respectively. The *sf3b1.L* and *S*, *ndufs1.L* and *S*, and *fn1.L* and *S* loci were mapped on the q arm of XBO Chr9_10L and 9_10S, respectively. Different distances from the centromeres in the *sf3b1*, *ndufs1*, and *fn1* signals are visible. A white line indicates the centromere position. For the female 8L chromosomes, the labeling of the sex chromosomes as W or Z is arbitrary because we are unable to distinguish them using cytogenetics.

To accomplish (ii) and (iii), we mapped five t(9;2) translocation-associated genes. According to the *X. borealis* genome database, *gyg2.L* probe mapped to XBO Chr2L spanning 34,388,249-34,388,569 bp, *cept1* to XBO Chr2S 52,504,791-52,505,068 bp, *sf3b1.L* to XBO Chr9_10L 93,569,359-93,569,596 bp, *ndufs1*.*L*(*S*) to XBO Chr9_10L(S) 89,927,074-89,927,354 (78,525,925-78,526,222) bp, and *fn1*.*L* to XBO Chr9_10L 82,455,252-82,455,624 bp. Because the *X. borealis* genome does not have identified centromere coordinates, it is not possible to determine chromosomal arms carrying the studied genes. Previous studies identified these genes in *X. tropicalis* and *X. pygmaeus*: *gyg2* and *cept1* genes on the short (p) arm; *sf3b1*, *ndufs1*, and *fn1* on the long (q) arm [44,61]. Our *gyg2*.*L* and *S* TSA probes mapped to the p arm of XBO Chr2L and 2S, the *cept1*.*L* probe mapped to the p arm of XBO Chr2L and 2S. The *sf3b1.L* and *S*, *ndusf1.L* and *S*, and *fn1.L* and *S* probes hybridized to the q arm of XBO Chr9_10L and 9_10S respectively (Fig. 4; Supplementary material, Figs S5-S9). According to the localization and order of the *sf3b1*, *ndusf1*, and *fn1* genes on the *X. borealis* Chr9_10L and 9_10S we can conclude that ancestral chromosomes 9 and 10 fused by their long arms as it happened in *X. laevis* ancestor. The translocation between chromosomes 9 and 2 did not happen in *X. borealis*.

### 3.5. Identification of *X. borealis* chromosomes based on p and q arms measurements and statistical analysis

All four individuals, regardless of sex, consistently had 36 chromosomes (2*n*c:=c:4*x*c:=c:36); for *X. borealis* this corresponds to *n* = 18, *x* = 9. This finding confirms that all analysed individuals are biological diploid yet evolutionary paleotetraploid, which coincides with Tymowska [27].

The arms of each chromosome were measured from 25 metaphase spreads (as a pool of 10 male and 15 female metaphases) and the median values of *l*, *r*_1_ and *i* were calculated (Table 1). The value of *l* was quantified as a percentage of the sum of *l* for all 36 chromosomes (sum of *l* per each chromosome is equal to 50) to account for variation in resolution and pixel size of our images. The karyotype of *X. borealis* consists of 10 pairs of metacentric (m), 5 pairs of submetacentric (sm), and 3 pairs of subtelocentric (st) chromosomes. Neither acrocentric chromosomes (*i* interval 0–12.5) nor telocentric chromosomes (no p arm; *i* = 0) were present in the *X. borealis* karyotype.

Subsequently, female (Fig. 5a,c,e) and male (Fig. 5b,d,f) metaphase spreads were analysed separately to identify sex-specific differences in chromosome morphology, specifically whether there is statistically significant difference in *l* or *i* between male and female chromosome 1L, 1S, etc. Each chromosome was plotted on a graph based on *i* (x axis) and *l* (%, y axis) to determine whether pairs of homoeologs had similar chromosomal morphology and whether they grouped closely together in females (Fig. 5a) and males (Fig. 5b). The *l* and *i* values corresponding to each chromosome 1L-9_10S were plotted as well (females Fig. 5c,e; males Fig. 5d,f).

**Figure 5.**
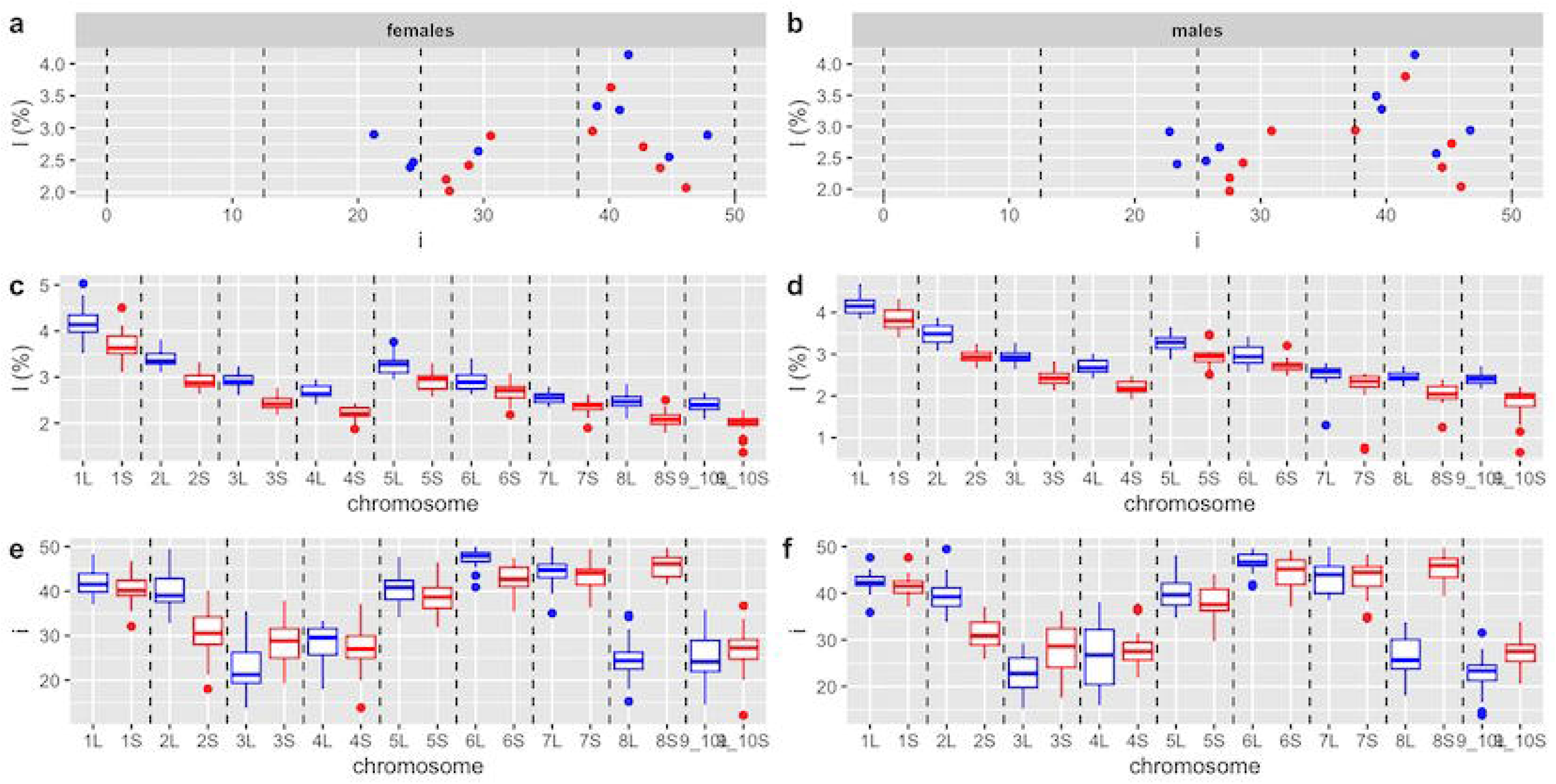
Morphometric analysis and statistics of *X. borealis* chromosomes. The L-subgenome is shown in blue and the S-subgenome in red. Panels a,b show the relationship between centromeric index (*i*), x-axis, and chromosome length (*l*), y-axis. Black dashed vertical lines delineate the intervals 0–12.5, 12.5–25, 25–37.5, and 37.5–50, corresponding to acrocentric, subtelocentric, submetacentric, and metacentric chromosomes, respectively. The plotted values of *i* and *l* are medians for each chromosome. Panels c,d show intrachromosomal variation of *l* value (y axis) for the haploid complement of 18 *X. borealis* female (c) and male (d) chromosomes (x axis). Panels e,f represent intrachromosomal variation of *i* value (y axis) for the haploid complement of 18 *X. borealis* female (e) and male (f) chromosomes (x axis). c,d,e,f: Black dashed vertical lines define pairs of homoeologous chromosomes. Upper and lower whiskers show minimum and maximum values, respectively; boxes involve the lower (*Q*_1_) and upper (*Q*_3_) quartiles; horizontal lines inside the boxes indicate the median values (*Q*_2_); outliers are indicated by blue (for L) and red (for S) points above and below the whiskers.

A measure of statistical dispersion, the interquartile range (*Q*_1_–*Q*_3_), was used to evaluate the extent of morphological divergence of each pair of homoeologous chromosomes. The highest divergence in *l* between homoeologous chromosomes was found between chromosomes 3L and 3S. The *Q*_1_–*Q*_3_ of chromosomes 3L and 3S ranged from 2.84 to 3.02% and from 2.34 to 2.53%, respectively (Fig. 5c,d; Supplementary material, Table S2). The largest difference in *i* between homoeologous chromosomes was between chromosomes 8L and 8S, which differed by 22.48–28.06 and 43.31–47.58 within *Q*_1_–*Q*_3_, respectively. Each of homoeologous chromosomes 8L and 8S was classified into different morphological categories (st, and m; Table 1). The second greatest variation based on the *Q*_1_–*Q*_3_c:*i* interval was between 2L (37.40–42.31) and 2S (28.16–34.10) (Fig. 5e,f; Supplementary material, Table S3). This homoeologous pair was also assigned into different categories (m, and sm; Table 1). A two-way ANOVA revealed statistically significant differences in *l* between all homoeologous pairs, and in *i* between all pairs except chromosomes 4L/4S and 7L/7S. A marginally significant effect of sex in *l* was observed for Chr9_10 (*p* = 0.05). Tukey’s test revealed one interaction on the level of the same subgenome and different sex for *l* values between male and female chromosomes 9_10S (borderline significance p < 0.05). All statistical significance is shown in Table 2. Analysis of the *l* value confirmed the appropriateness of the L (long) and S (short) nomenclature, as statistically significant differences in length exhibit between homoeologs of every pair.

**Table 2.**
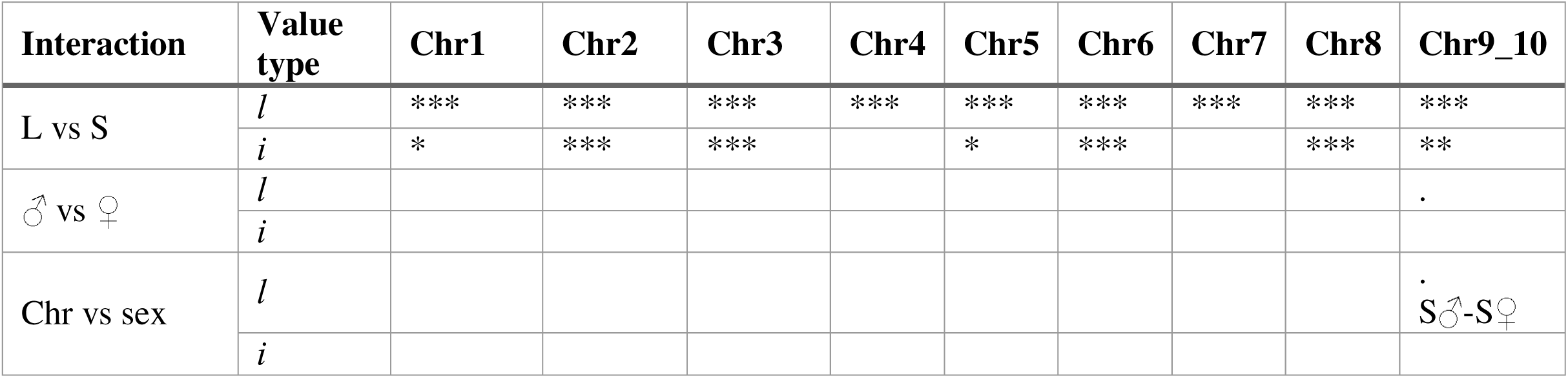
Analysis of statistical significance of measured *l* and *i* values of individual chromosomes (L vs S), sex (♂ vs ♀), and interaction of chromosomes and sex (Chr vs sex). Each statistically significant interaction is defined by either L or S chromosomes and either males or females. Difference between male and female *l* values within the same chromosome was found in chromosome 9_10S. Significantly different chromosomes are depicted by significance codes ***, **, and * showing *p* values of *p*c:<c:0.001, *p*c:<c:0.01, and *p*c:<c:0.05, respectively.

### 3.6. Whole-genome mapping

Four synteny plots across three *Xenopus* species and their subgenomes (*X. tropicalis*, *X. laevis* L and S, *X. borealis* S and L) were generated to investigate chromosome rearrangements (Fig. 6a; Supplementary material, Fig. S10) and their timing (Fig. 6b). Each chromosome was dissected from C-banded metaphase spread (Fig. 6c) and inserted to the synteny lines to visualize rearranged chromosomal segments and position of centromeres. A fusion between ancestral *X. borealis* and *X. laevis* chromosomes 9 and 10 is clearly visible. Chromosome telomeres fused with their q arms. A large inversion is present on XLA 3S and XBO 3S indicating that the inversion occurred in a common diploid S-ancestor of *X. borealis* and *X. laevis* or shortly after polyploidization common for *X. borealis* and *X. laevis*. This inversion spans almost the entire length of the chromosome except telomeres. Inversions on chromosome 2L, 5L common for both *X. borealis* and *X. laevis* occurred most likely in the L-ancestor before tetraploidization. Given the additional inversion on XBO 2L smaller than inversion on XLA 2L indicates that after the inversion most recent L-ancestor of *X. borealis* and *X. laevis*, the portion (approximately half) of the inverted region was re-inverted recently in the *X. borealis* L-subgenome. Inversion on the p arm of chromosomes XLA 2S and 4S are not present on XBO 2S and 4S, respectively, which shows that these inversions happened very recently in *X. laevis* S subgenome after divergence of *X. borealis* and *X. laevis* lineages, most likely after tetraploidization. Multiple small inversions are detected on *X. borealis* chromosomes 1L, 1S, 2L, 2S, 3L, 7S, 8L, showing that these inversions have independent and recent origins in the lineage forming *X. borealis* species. Another small inversion is visible on the distal part of the p arm of chromosomes 1L and 1S in both *X. borealis* and *X. laevis*. The inversion is evolutionarily old and has similar timing and as fusion between ancestral chromosomes 9 and 10. A translocation was identified within XLA 8S. This locus moved from pericentromeric to telomeric region of the same chromosome and subsequently inverted. The translocation did not occur in XBO 8S, indicating its recent origin in *X. laevis* lineage.

**Figure 6.**
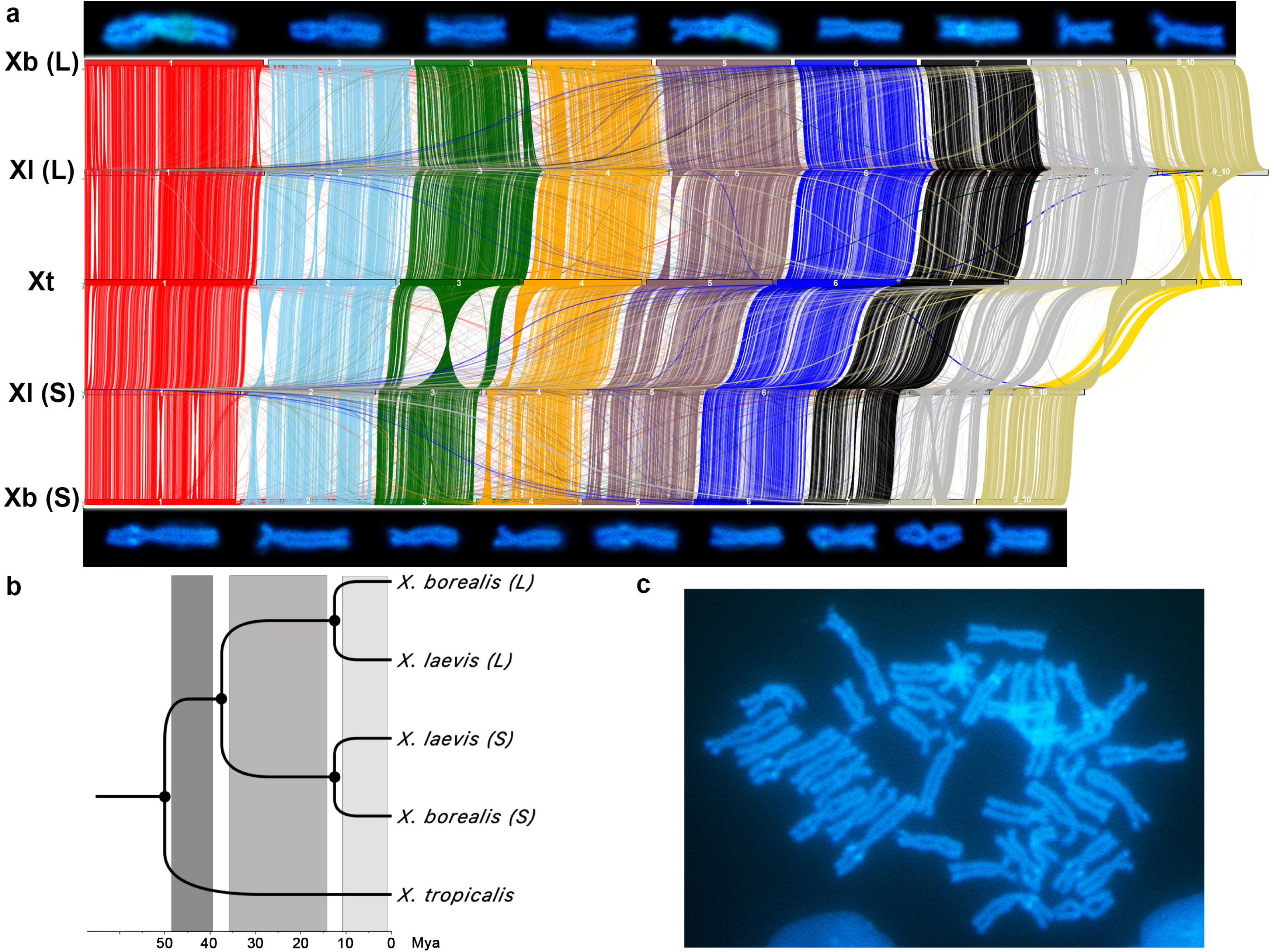
Whole-genome synteny analysis. (a) The *X. tropicalis* (Xt) genome containing 10 haploid chromosomes (middle line) aligned with *X. laevis* L- (Xl (L)) and S-subgenomes (Xl (S)) (lines above and below *X. tropicalis*). The *X. laevis* L- and S-subgenomes mapped to the *X. borealis* L- (Xb (L)) and S-subgenomes (Xb (S)) (top and bottom lines). Each haploid chromosome is graphically depicted above or below the synteny map. (b) Phylogenetic tree involving two studied tetraploids and *X. tropicalis* as reference species with estimated timescale. The tree was estimated based on the *rag1* sequence using the Neighbor-Jioining method and the Tamura-Nei model, implemented in Geneious Prime (v2026.0.2.). Divergence of subgenera *Xenopus* and *Silurana* represents the deepest node approximately 50 Mya, followed by split of the L- and S-subgenomes 30-35 Mya. Ancestral, intermediate, and recent temporal strata of rearrangements are highlighted with dark, medium, and light gray, respectively. Nodes and divergence points serve as delineating boundaries and are not assigned to specific temporal strata. (c) C-banded metaphase plate, from which chromosomes were cut and inserted to the synteny map.

Using the whole-genome synteny analysis, we categorized rearrangements into three temporal strata. (i) Ancestral rearrangements that occurred before both subgenome divergence and polyploidization, i.e., the fusion of ancestral chromosomes 9 and 10 and small distal inversions on the p arm of 1L and 1S. (ii) Intermediate subgenome-specific rearrangements, occurring in either L- or S-subgenome in both *X. borealis* and *X. laevis* (inversions 2L, 3S, and 5L).

These events occurred after the divergence of the diploid progenitors but prior to the speciation of the *borealis-laevis* clade. However, whether they occurred before or after the polyploidization event remains to be elucidated. (iii) Recent subgenome- and species-specific rearrangements, occurring post-speciation in individual lineages (e.g., inversions on xla 2S, 4S, 8S, xbo 1S, 1L, 2L, 2S, 3L, 7S, 8L, as well as the xla 8S translocation).

In total, we identified 16 intra-chromosomal rearrangements and confirmed the ancestral fusion of chromosomes 9 and 10 in the evolution of *X. borealis* and *X. laevis*. Rearrangements occur continuously in three temporal strata throughout both subgenomes during approximately 50 My of *Xenopus* evolution.

## 4. Discussion

High-resolution whole-genome sequencing combined with molecular cytogenetics permit the evaluation of the changes in genome structure over time and across species. In this study, we employed these methods to investigate structural evolutionary landscape in the African clawed frog genus *Xenopus*, using two tetraploids *X. borealis* and *X. laevis* alongside with the diploid *X. tropicalis*.

Our characterization of the *X. borealis* karyotype (2*n* = 4*x* = 36) reveals that its sex chromosomes remain cytologically homomorphic. Despite the application of chromosome banding and mapping techniques, no heteromorphic patterns were identified, suggesting a lack of large-scale structural differentiation between the sex-determining pair. A karyotype of *Xenopus borealis* was first reported by Tymowska [27] but chromosome arms were not measured. Moreover, Tymowska [27] numbered chromosomes based on chromosome categories, while today-karyotype nomenclature is based on homology of each chromosome to *X. tropicalis* ortholog (sensu [30,47]), where the largest *X. borealis* chromosomes 1L and 1S are homologous to *X. tropicalis* chromosome 1, and *X. borealis* chromosomes 9_10L and 9_10S are homologous to *X. tropicalis* chromosomes 9 and 10. As previously revealed in related species from subgenus *Xenopus* [36,44], most of homoeologous chromosomes significantly differ in length and centromeric index (the L vs S counterparts) but are very similar within the submetacentric categories, such as if we compare 3L, 4L, 8L, and 9_10L. For better chromosome identification and detecting chromosome-specific markers, we implemented C-banding and FISH with rDNA and snDNA probes. Using these cytogenetic markers, we identified *X. borealis* chromosomes 1S, 5L, and 8L. Chromosome 8L is a sex chromosome and thus its marker for identification can be valuable for studies related to sex determination and its mechanisms. Using whole-chromosome painting probes from *X. tropicalis*, we identified homoeologous groups of X. borealis chromosomes 1, 6, 7, 8, 9, and 10.

Beyond basic characterization, molecular cytogenetics provides an insight into complex evolutionary processes such as allotetraploidy and asymmetric subgenome evolution [27,28,31,35,36,44]. Our findings are entirely consistent with an allotetraploid origin for both *X. borealis* and *X. laevis*. While early cytogenetic studies first proposed allotetraploidy as a primary driver of *Xenopus* evolution, the hybridization of two distinct ancestral progenitors approximately 17 Mya was later definitively confirmed by the subgenome-specific activation of DNA transposons [30,42,65].

### 4.1. Evolution-by-rearrangement

While traditional cytogenetics suggests that *Xenopus* has a conserved karyotype architecture, our in-depth gene-level investigation uncovered a complex landscape of structural changes that we grouped into three temporal strata. (i) Ancestral rearrangements that predate the subgenus radiation and are shared by the subgenomes of both *X. borealis* and *X. laevis*, but are absent in *X. tropicalis*. Key examples include the fusion of chromosomes 9 and 10 and distal p-arm inversions on chromosomes 1L and 1S. This fusion is corroborated by cross-species whole-chromosome painting and high-resolution sequencing [30,40,41,44,55], with an estimated timing between 35 and 50 Mya (Fig. 6b). (ii) Intermediate rearrangements that occurred in either one subgenome in both tetraploids. These rearrangements occurred after the divergence of the diploid progenitors but prior to the speciation of *X. borealis* and *X. laevis* approximately 15-35 Mya (inversions 2L, 3S, and 5L; Fig. 6a,b). These correspond to the "L/S common inversions" recently identified by Suda et al. [42]. (iii) Recent rearrangements are subgenome-, lineage- and species-specific events that occurred after speciation roughly 15 Mya or even more recently. Examples include inversions on *X. laevis* 2S, 4S, 8S, *X. borealis* 1S, 1L, 2L, 2S, 3L, 7S, 8L, as well as the *X. laevis* 8S translocation (Fig. 6a,b).

Our investigation identified a similar frequency of rearrangements across both subgenomes, which aligns with recent findings by Almojil et al. [66]. Notably, the prevailing paradigm that the L-subgenome is inherently more structurally stable than the S-subgenome remains supported by significant evidence from divergent gene expression, differentiated transposable element content, pseudogenization [30,33,42]. Consequently, we do not disprove the model of asymmetric subgenome evolution; rather, our results suggest that structural rearrangements may follow a more symmetric evolutionary trajectory than previously hypothesized.

While we localized a telomeric inversion to *X. laevis* chromosome 5L, Session et al [30] identified the same inversion on chromosome 5S. This contradiction suggests a need for refined assembly and genome mapping in *Xenopus* polyploid models. Because this inversion is conserved across both studied tetraploids, we classified it within the intermediate temporal stratum.

The sex chromosomes of *X. borealis* are female-heterogametic (ZZ males, ZW females) with yet an unidentified sex-determining gene localized to chromosome 8L [64]. In this study, we tested the hypothesis that a putative inversion within the sex-determining locus of the female-specific W chromosome suppresses recombination between the Z and W chromosomes and facilitates structural divergence. We physically mapped three single-copy genes (*sf-1.L*, *ar.L*, and *sox3.L*) located within the region of suppressed recombination [64] to evaluate whether their position and linear order are identical on the Z and W chromosomes. Our cytogenetic mapping results correspond with locations in the *X. borealis* male reference genome [41], with *sf-1.L* and *ar.L* localized to the telomere and pericentromere of the p arm, respectively, and *sox3.L* to the q arm (Fig. 4). Comparative analysis of a female specimen revealed that the gene order and centromeric indices are the same on the Z and W chromosomes. Thus, we found no evidence of large-scale structural divergence or macro-inversions and no indication of the suppressed recombination between sex chromosomes due to inversion, which is a proposed mechanism in other vertebrate taxa [67,68]. While our results suggest a lack of large-scale chromosomal reorganization, we cannot exclude the possibility of micro-inversions or rearrangements involving different loci than targeted by our FISH probes. Thus far, the *X. borealis* W chromosome remains cytologically homomorphic, distinguished from the Z only by sex-specific SNPs [64].

Another rearrangement that we explored was the translocation from chromosome 9 to 2 observed in *X. mellotropicalis*. Although whole-chromosome painting probes from *X. tropicalis* chromosome 2 failed to detect any signals and whole-chromosome painting probe 9 showed only faint hybridization on the q arm, single-copy gene mapping of t(9;2) translocation-associated genes did not identify the translocation in *X. borealis*, showing the translocation to be a *Silurana*-specific event as revealed by Knytl et al. [34] and [35].

### 4.2. Tandem repeat loci

Across the Pipidae, NORs have been mapped using multiple approaches (FISH, C-banding, quinacrine mustard fluorescence), consistently identifying a single locus (one homologous pair) but with significant interspecific spatial variation, i.e., on distinct chromosomes and positions [27,34,35,44,52,69]. This positional variability is attributed to a jumping mechanism [44,70], originally described for the movement of transposable elements [71]. For the *Silurana* subgenus, the NOR location is conserved on chromosome 7q, likely representing the ancestral state for the entire genus *Xenopus*. In contrast, subgenus *Xenopus* exhibits diverse localizations across both subgenomes, including chromosomes 3L, 5L, and 6S [27,28,44,52]. The NOR on chromosome 6S is hypothesized to jump from the L-subgenome [44].

A notable discrepancy emerged regarding the NOR mapping in *X. borealis*. While BLAST analysis of our rDNA probe identified multiple copies on chromosomes 4L of the male reference assembly [41], our cytogenetic mapping primarily detected the signal on 5L, with an additional lower-abundance signal on 4S. Although inter-individual variation in NOR location has not been previously reported in Pipidae, the missmapping error may stem from assembly artifacts in the initial *X. borealis* reference genome. Interestingly, *X. muelleri*, a sister species to *X. borealis*, also harbors NOR on chromosome 5L, but it is the pericentromeric region of the q arm, which is a non-homologous position relative to the 5Lp telomeric signal observed in *X. borealis*. These data confirm that rDNA mobilization has altered NOR positions in either the *X. muelleri* or *X. borealis*, though the ancestral state for the subgenus *Xenopus* and the root cause of the current genomic-cytogenetic inconsistency remain to be resolved.

The genomic distribution of snDNA repeats across Pipidae is relatively stable, with U1 localized on chromosome 1 and U2 on chromosome 8. However, copy numbers per these loci vary in individual subgenomes throughout the reduction-expansion mechanism (sensu [36], see also[26]). Subgenome-specific localizations vary among species; for example, U1 signals are restricted to chromosome 1L in *X. pygmaeus* but to 1S in *X. laevis*, while species such as *X. muelleri* and tetraploid *Silurana* maintain signals on both homoeologs. A similar pattern exists for U2, with FISH signals on 8L (*X. laevis*), 8S (*X. pygmaeus, X. allofraseri*), or both homoeologs (*X. muelleri, Silurana*). *Xenopus borealis* represents a unique exception, in addition to the 8S localization suggested by the genome assembly, our cytogenetic mapping identified a primary signal on 8L and an additional, low-abundance signal on 4S. Our physical mapping of tandem repeats can serve as a marker for identification *X. borealis* chromosomes 1S (U1 snDNA), 5L (NOR), 8L (U2 snDNA).

The U1 and U2 snDNA repeats undergo an evolutionary trajectory distinct from the categorical deletion of the NOR. While BLAST analyses identified several copies of U1 and U2 snDNAs on *X. borealis* chromosomes 1S and 8S, respectively. The copy number per the U2 locus is markedly reduced in the S-subgenome, compared to the L-subgenome and the diploid *X. tropicalis* genome. Conversely, the copy number per the U1 locus was expanded in the S-subgenome as evidenced by FISH. The reduction-expansion dynamics in tandem-repeat copy numbers is consistent with recent findings in *X. laevis* [36], further highlighting the asymmetric evolution of repetitive elements as a hallmark of subgenome divergence.

## 5. Conclusions

Our findings suggest that different classes of genomic variation follow distinct evolutionary trajectories in polyploid *Xenopus*. While the S-subgenome is characterized by a significantly higher number of deletions [30]—as exemplified by the functional loss of NORs observed here and in previous studies [34,35,44]—this asymmetry appears most prominent in the early stages of *Xenopus* diversification—after the divergence of two diploid ancestors but before speciation of *X. laevis* and *X. borealis* [30]. In contrast, identified structural rearrangements (inversions, translocation, and fusion) do not exhibit the same subgenome bias. Instead, we found the balanced distribution of these events across both the L- and S-subgenomes in three temporal evolutionary strata through proportional accumulation during 50 My of evolutionary history of subgenus *Xenopus*. Moreover, we did not detect any inversions on the *X. borealis* W chromosome. This finding challenges the traditional hypothesis that large-scale inversions suppress recombination, even though the eastern population is known to have a large (54.1 Mb) non-recombining region between the Z and W chromosomes [41,72]. Therefore, a mechanism other than large inversions is responsible for maintaining this such extensive sex-specific locus within the highly homomorphic sex chromosomes of *X. borealis*.

## Supplementary material

**Table S1:** Species name, markers used in FISH analyses, organs used for DNA and RNA extractions, PCR primer sequences, lengths of amplicons, working designation of plasmid clones selected for FISH, GenBank accession numbers.

**Table S2**: Minimum (*Q*_1_) and maximum (*Q*_3_) values, and interquartile range (IQR) of *l* values for each individual chromosome of *Xenopus borealis*.

**Table S3**: Minimum (*Q*_1_) and maximum (*Q*_3_) values, and IQR of *i* values for each individual chromosome of *X. borealis*.

**Figure S1: FISH with 5S ribosomal probe in Black & White mode.** (a) DAPI shows intense spots on telomeres of some subtelocentric chromosomes. (b) 5S rDNA FISH shows intense spots on telomeres of the other chromosomes than DAPI does.

**Figure S2: FISH-TSA with positive *sox3.L* signals on the *X. borealis* metaphase spreads.** (a, b) Shows female metaphase spreads. (c, d) Shows male metaphase spreads. For both sexes the *sox3.L* gene (red) was localized on the q arm of *X. borealis* Chr8L (XBO Chr8L). Chromosomes were counterstained with DAPI (blue-green). Scale bars represent 10 μm.

**Figure S3: FISH-TSA with positive *ar.L* signals on the *X. borealis* metaphase spreads**. (a, b) Shows female metaphase spreads. (c, d) Shows male metaphase spreads. For both sexes the *ar.L* gene (red) was localized on the p arm of XBO Chr8L, respectively. Chromosomes were counterstained with DAPI (blue-green). Scale bars represent 10 μm.

**Figure S4: FISH-TSA with positive *sf-1.L* signals on the *X. borealis* metaphase spreads.** (a, b) Shows FISH-TSA on female metaphase spreads. (c) Shows FISH-TSA (red) and U2 snDNA (green) on female metaphase spreads. Both genes *sf-1.L* and U2 snDNA were localized on XBO Chr8L. (d) Shows FISH-TSA on male metaphase spreads. For both sexes the *sf-1.L* gene was localized on the p arm of XBO Chr8L, respectively. Chromosomes were counterstained with DAPI (blue-green). Scale bars represent 10 μm.

**Figure S5: FISH-TSA with positive *gyg2* signals on the *X. borealis* metaphase spreads.** The *gyg2.L* and *S* genes (red) were localized on the p arm of XBO Chr2L and 2S, respectively. Chromosomes were counterstained with DAPI (blue-green). Scale bars represent 10 μm.

**Figure S6: FISH-TSA with positive *cept1* signals on the *X. borealis* metaphase spread.** The *cept1.L* and *S* genes (red) were localized on the p arm of XBO 2L and 2S, respectively. The *cept1.S* probe hybridized to a single Chr2S gametologue as shown on the metaphase spread on the right side of the panel. Chromosomes were counterstained with DAPI (blue-green). Scale bars represent 10 μm.

**Figure S7: FISH-TSA with positive *sf3b1* signals on the *X. borealis* metaphase spreads.** The *sf3b1.L* and *S* genes (red) were localized on the q arm of XBO Chr9_10L and 9_10S, respectively. Chromosomes were counterstained with DAPI (blue-green). Scale bars represent 10 μm.

**Figure S8: FISH-TSA with positive *ndufs1* signals on the *X. borealis* metaphase spreads.** The *ndufs1.L* and *S* genes (red) were localized on the q arm of XBO Chr9_10L and 9_10S, respectively. Chromosomes were counterstained with DAPI (blue-green). Scale bars represent 10 μm.

**Figure S9: FISH-TSA with positive *fn1* signals on the *X. borealis* metaphase spreads.** The *fn1.L* and *S* genes (red) were localized on the q arm of XBO Chr9_10L and 9_10S, respectively. Chromosomes were counterstained with DAPI (blue-green). Scale bars represent 10 μm.

**Figure S10: Whole-genome synteny analysis.** (a) The *X. tropicalis* (Xt) genome containing 10 haploid chromosomes (middle line) aligned with the *X. borealis* L-subgenome (xb L; top) and S-subgenome (xb S; bottom). (b) Similar alignment of the *X. tropicalis* (Xt) genome (middle) with the *X. laevis* L-subgenome (xl L; top) and S-subgenome (xl S; bottom). In both species, the *X. tropicalis* chromosomes serve as the reference for subgenome comparisons.

## Data statement

All experimental data supporting the findings of this study are available within the article and its supplementary materials. DNA sequences have been deposited in GenBank (accession numbers are provided in Supplementary Table S1). Data analysis pipeline and associated files are available on GitHub at https://github.com/martinknytl (see Materials and Methods, sections 2.5 and 2.6 for details).

## Acknowledgements

We thank Sarah Quigley and Marta Farre Belmonte for nice help with synteny maps generation and its related troubleshooting. Also, we thank XXX anonymous reviewers for their constructive comments on early version of the manuscript.

## Declaration of generative AI use

During the preparation of this work, the authors used Google AI Studio (Gemini) to improve the language, grammar, and stylistic clarity of the manuscript. Following the use of this tool, the authors reviewed and edited the generated content and take full responsibility for the integrity and accuracy of the final version.

## Funding

The research was supported by the Grant Agency of Charles University (GAUK), project 186024 [BB] and [MK], the Cooperatio Program in Biology at Charles University [TT, VK], the Natural Science and Engineering Research Council of Canada (RGPIN-2024-05290) [BJE], the P JAC project CZ.02.01.01/00/22_010/0002902 MSCA Fellowships CZ—UK [MK]. Open access publishing supported by the institutions participating in the CzechELib Transformative Agreement.

## Author contributions: CRediT

Conceptualization: [BJE] and [MK]; Data curation: [MK]; Formal analysis: [MK]; Funding acquisition: [BB], [TT], [VK], [BJE] and [MK]; Investigation: [BB], [NRF], [TT], [JV], [MP], [HČ], [SK] and [MK]; Methodology: [MK]; Project administration: [MK]; Resources: [BJE]; Software: [BJE] and [MK]; Supervision: [MK]; Validation: [MK]; Visualization: [MK]; Writing – original draft: [MK]; Writing – review and editing: [BB], [VK], [BJE] and [MK].

## Research Ethics Statement

Charles University has registered experimental breeding facilities for pipid frogs (16OZ12891/2018-17214, 37428/2019-MZE-18134). All experimental procedures involving frogs were approved by the Institutional Animal Care and Use Committee of Charles University, according to the directives from the State Veterinary Administration of the Czech Republic, reference number MSMT-20585/2022-4 issued by the Ministry of Education, Youth and Sport of the Czech Republic (MK is a manager of the experimental project on living Xenopus animals). MK is a holder of the Certificate of professional competence to design experiments according to §15d(3) of the Czech Republic Act No. 246/1992 coll. on the Protection of Animals against Cruelty (Registration number CZ 03973), provided by the Ministry of Agriculture of the Czech Republic.

## Notes

### Competing Interest Statement

The authors have declared no competing interest.

