## Supplementary Materials for "Reconstructing 50 million years of *Xenopus borealis* evolution: three temporal strata of DNA rearrangements and persistent sex chromosome homomorphism"

**Table S1**: Species name, markers used in FISH analyses, organs used for DNA and RNA extractions, PCR primer sequences, lengths of amplicons, working designation of plasmid clones selected for FISH, GenBank accession numbers.

| Species | Gene symbol | Name of the gene | Tissue | Type of sequence | Primer sequence | Size (bp) | Clone | NCBI number |
|---|---|---|---|---|---|---|---|---|
| ***Xenopus borealis*** | *fn1* | *fibronectin* | spleen | F1 | 5′-AATGGGAGCGCACCTACCT-3′ | 1329 | S: A | |
| | | | | R1 | 5′-CATCCACAATGCACTGGTCT-3′ | | L: F | |
| | *sf3b1* | *splicing factor 3b subunit 1* | | F | 5′-TTGCGAAAACACACGAAGATATTG-3′ | 1410 | S: C | |
| | | | | R1 | 5′-GTGTAGATTCATCAACATCAACCA-3´ | | L: B | |
| | *ndufs1* | *NADH:ubiquinone oxidoreductase core subunit S1* | | F | 5′-AGAAACCCATTGTGGTGGTTG-3′ | 934 | S: J | |
| | | | | R2 | 5´-GGTAATAAACCATTTTAATAAGTTTACAAG-3´ | | L: C | |
| | *cept1* | *choline/ethanolamine phosphotransferase 1* | | F | 5′-GTGGAAATCCCTACCAAACAG-3′ | 766 | H | |
| | | | | R2 | 5´-CACTTAGTGATGATTAAAACGTGC-3´ | | | |
| | *gyg2* | *glycogenin 2* | | F | 5′-TACTGTCAAGGAGCCCTGG-3′ | 1179 | S: L | |
| | | | | R2 | 5′-GCAAAACTAAATAGCTCTAATTTATTTG-3´ | | L: A | |
| | *ar* | *androgen receptor* | liver | F1 | 5′-CCTGCTGCCCATTGATTATTAC-3´ | 1219 | F | |
| | | | | R1 | 5′-ATACACAGCCAGGGTGATGC-3´ | | | |
| | *sf-1* | *nuclear receptor subfamily 5 group A member 1* | testes | F1_F5 | 5′-AAGTACCGCATGCTCTTGGT-3′ | 1430 | A | |
| | | | | R | 5′-TAGCAAAGGGGCAGAGAAAA-3′ | | | |
| | *sox3* | *SRY-box transcription factor 9* | brain | F1_F5 | 5′-CCGCACATCTCTTTTTGTCA-3′ | 1110 | B | |
| | | | | R | 5′-CACAACTCGGAGATCAGCAA-3′ | | | |

**Table S2**: Minimum ($Q_1$) and maximum ($Q_3$) values, and interquartile range (IQR) of *l* values for each individual chromosome of *Xenopus borealis*.

| Chromosome | $Q_1$ | $Q_3$ | IQR |
|---|---|---|---|
| 1L | 3.985546 | 4.301331 | 0.3157841 |
| 1S | 3.569395 | 3.935485 | 0.3660898 |
| 2L | 3.284347 | 3.583425 | 0.2990776 |
| 2S | 2.810655 | 3.037002 | 0.2263475 |
| 3L | 2.841283 | 3.023852 | 0.1825693 |
| 3S | 2.335110 | 2.532317 | 0.1972061 |
| 4L | 2.587145 | 2.813891 | 0.2267464 |
| 4S | 2.102166 | 2.334906 | 0.2327407 |
| 5L | 3.095836 | 3.389471 | 0.2936346 |
| 5S | 2.790306 | 3.021870 | 0.2315640 |
| 6L | 2.775153 | 3.072741 | 0.2975885 |
| 6S | 2.558139 | 2.786656 | 0.2285173 |
| 7L | 2.453435 | 2.644145 | 0.1907100 |
| 7S | 2.259295 | 2.447913 | 0.1886183 |
| 8L | 2.375513 | 2.552026 | 0.1765135 |
| 8S | 1.963730 | 2.193439 | 0.2297086 |
| 9_10L | 2.305364 | 2.516731 | 0.2113664 |
| 9_10S | 1.892011 | 2.080417 | 0.1884060 |

**Table S3**: Minimum ($Q_1$) and maximum ($Q_3$) values, and IQR of *i* values for each individual chromosome of *X. borealis*.

| Chromosome | $Q_1$ | $Q_3$ | IQR |
|---|---|---|---|
| 1L | 40.52455 | 43.82022 | 3.295670 |
| 1S | 39.14038 | 42.52874 | 3.388357 |
| 2L | 37.39512 | 42.31443 | 4.919315 |
| 2S | 28.15582 | 34.10160 | 5.945778 |
| 3L | 19.35484 | 26.34979 | 6.994949 |
| 3S | 24.82643 | 31.90403 | 7.077602 |
| 4L | 22.41230 | 31.92657 | 9.514274 |
| 4S | 25.21319 | 29.57746 | 4.364280 |
| 5L | 37.59705 | 42.36167 | 4.764618 |
| 5S | 36.25060 | 40.82840 | 4.577803 |
| 6L | 46.52406 | 48.64950 | 2.125435 |
| 6S | 41.56602 | 45.87979 | 4.313770 |
| 7L | 41.60547 | 45.88506 | 4.279593 |
| 7S | 41.26354 | 45.50287 | 4.239328 |
| 8L | 22.48062 | 28.05755 | 5.576934 |
| 8S | 43.30672 | 47.57509 | 4.268373 |
| 9_10L | 21.39207 | 26.46501 | 5.072944 |
| 9_10S | 24.86096 | 29.07801 | 4.217052 |

**Figure S1: FISH with 5S ribosomal probe in Black & White mode.** (a) DAPI shows intense spots on telomeres of some subtelocentric chromosomes. (b) 5S rDNA FISH shows intense spots on telomeres of the other chromosomes than DAPI does.

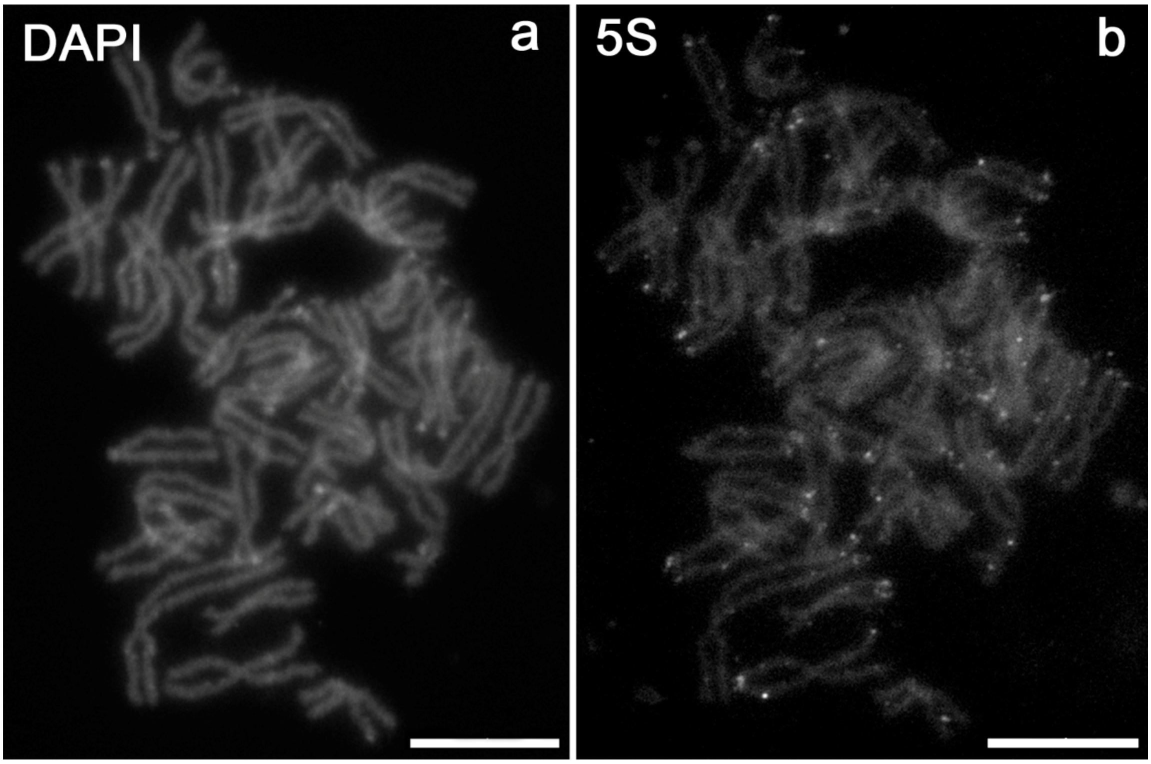

**Figure S2: FISH-TSA with positive *sox3.L* signals on the *X. borealis* metaphase spreads.** (a, b) Shows female metaphase spreads. (c, d) Shows male metaphase spreads. For both sexes the *sox3.L* gene (red) was localized on the q arm of *X. borealis* Chr8L (XBO Chr8L). Chromosomes were counterstained with DAPI (blue-green). Scale bars represent 10 µm.

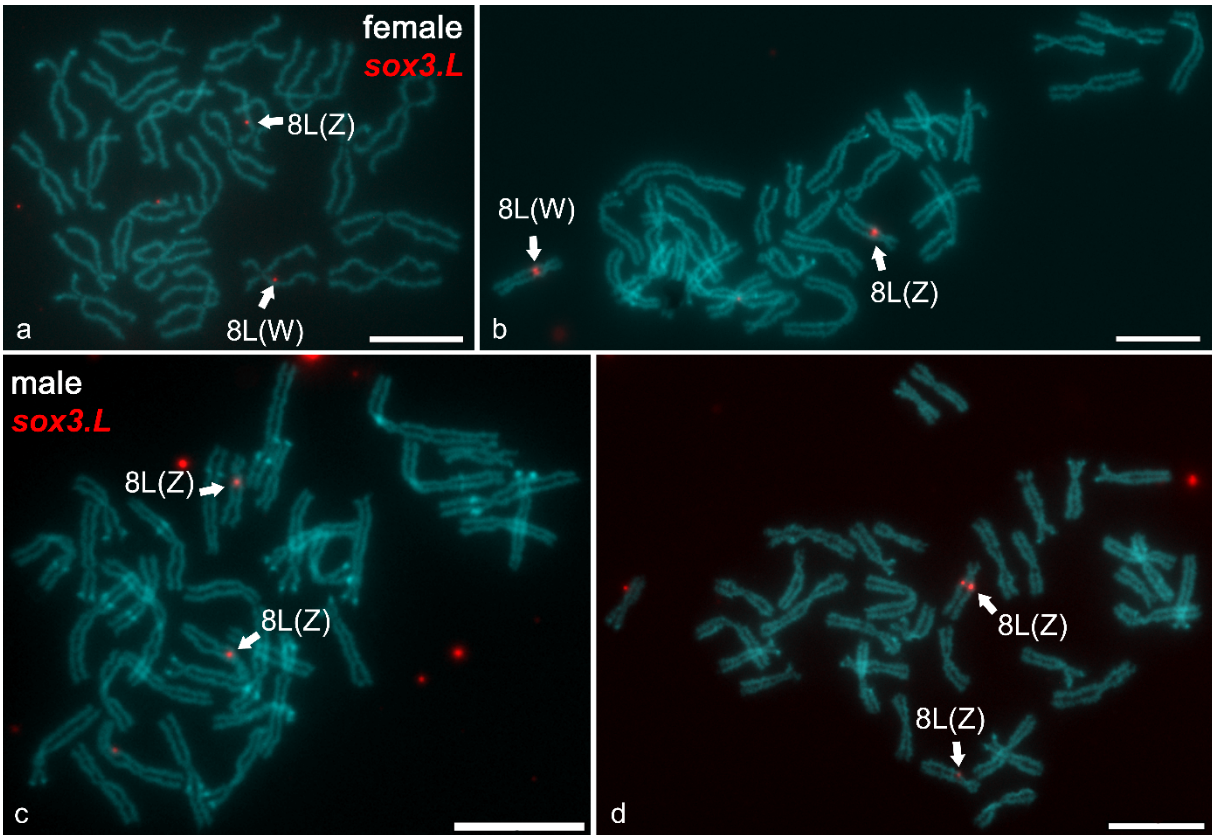

**Figure S3: FISH-TSA with positive *ar.L* signals on the *X. borealis* metaphase spreads**. (a, b) Shows female metaphase spreads. (c, d) Shows male metaphase spreads. For both sexes the *ar.L* gene (red) was localized on the p arm of XBO Chr8L, respectively. Chromosomes were counterstained with DAPI (blue-green). Scale bars represent 10 µm.

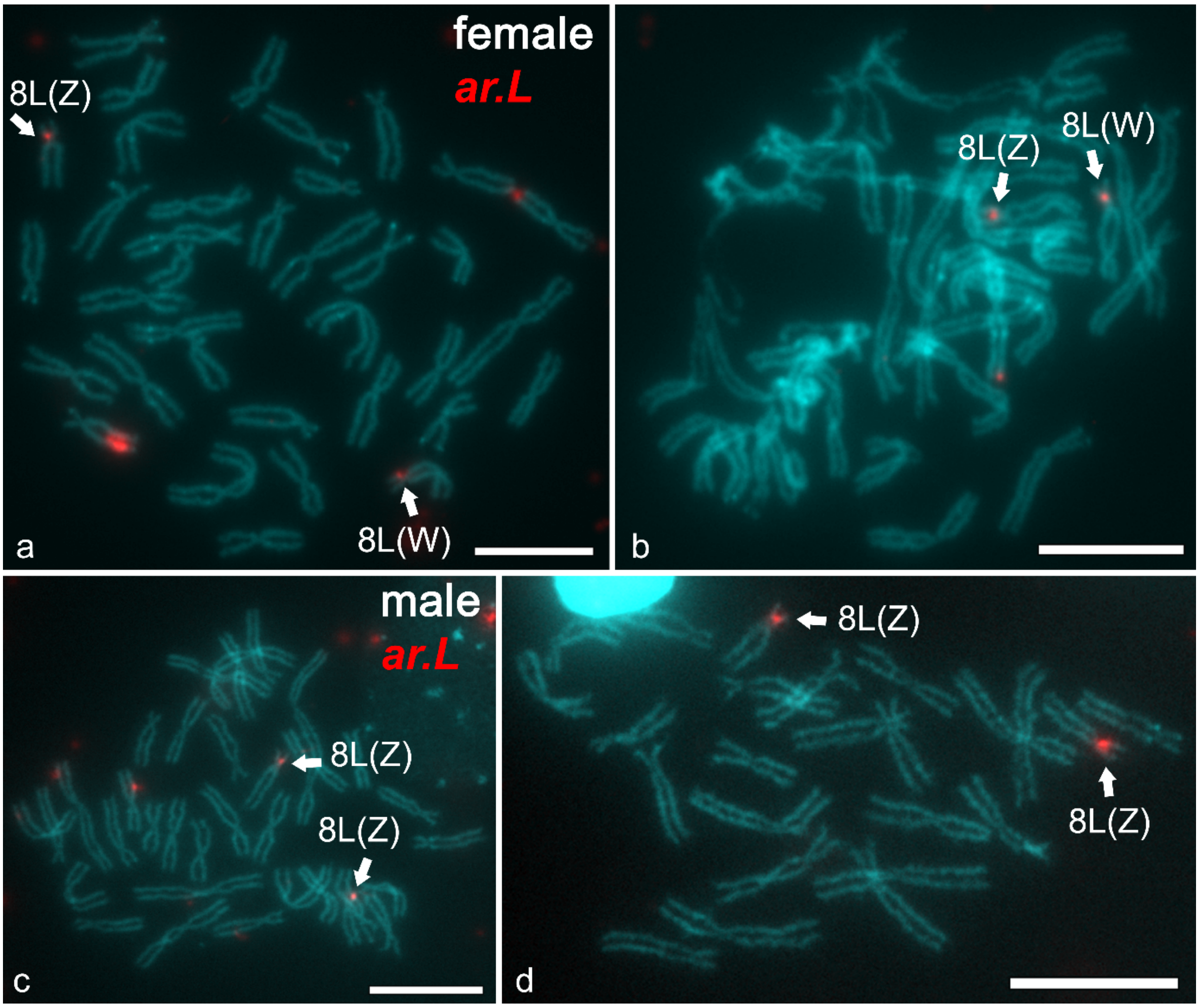

**Figure S4: FISH-TSA with positive *sf-1.L* signals on the *X. borealis* metaphase spreads.** (a, b) Shows FISH-TSA on female metaphase spreads. (c) Shows FISH-TSA (red) and U2 snDNA (green) on female metaphase spreads. Both genes *sf-1.L* and U2 snDNA were localized on XBO Chr8L. (d) Shows FISH-TSA on male metaphase spreads. For both sexes the *sf-1.L* gene was localized on the p arm of XBO Chr8L, respectively. Chromosomes were counterstained with DAPI (blue-green). Scale bars represent 10 µm.

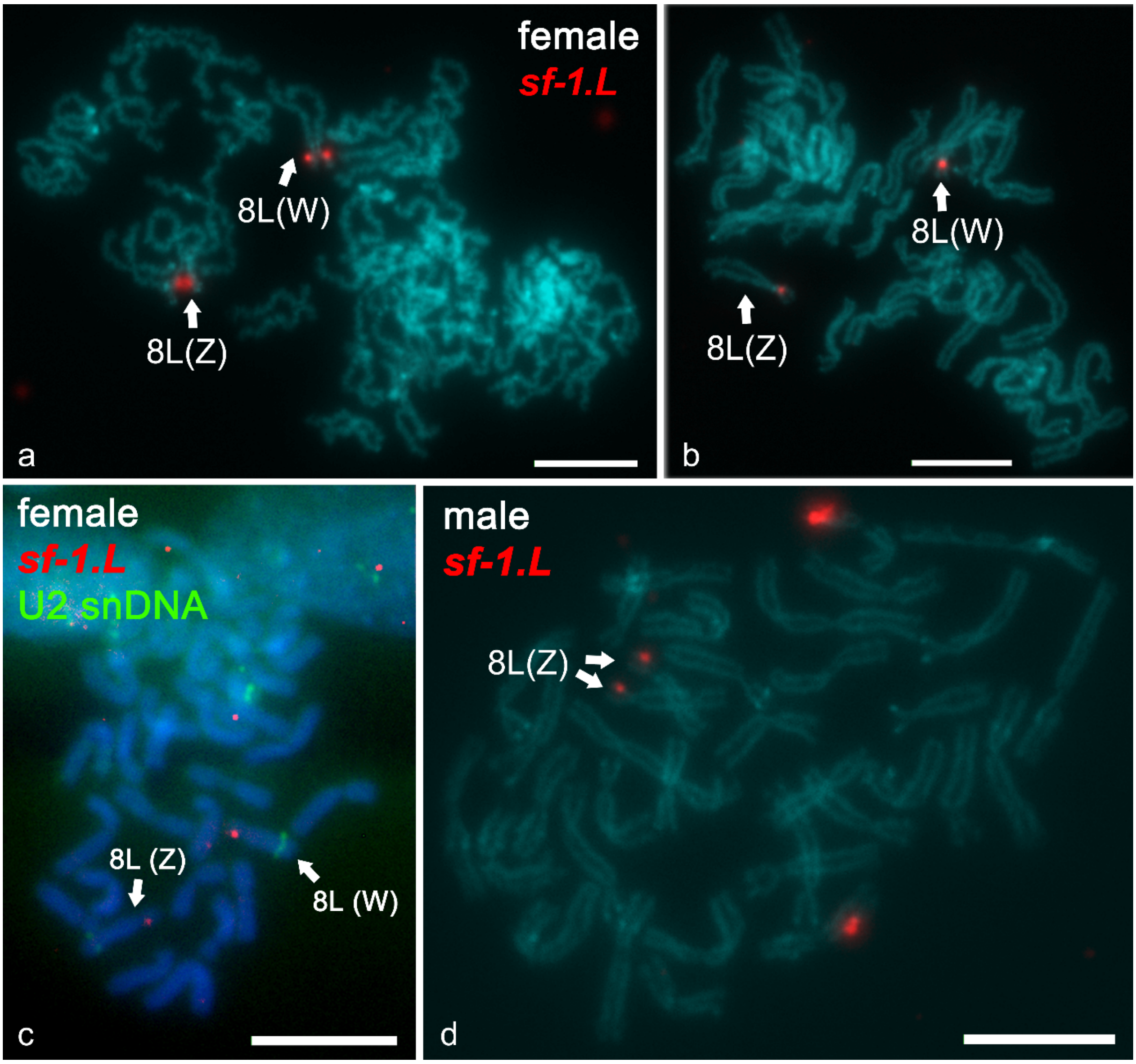

**Figure S5: FISH-TSA with positive *gyg2* signals on the *X. borealis* metaphase spreads.** The *gyg2.L* and *S* genes (red) were localized on the p arm of XBO Chr2L and 2S, respectively. Chromosomes were counterstained with DAPI (blue-green). Scale bars represent 10 µm.

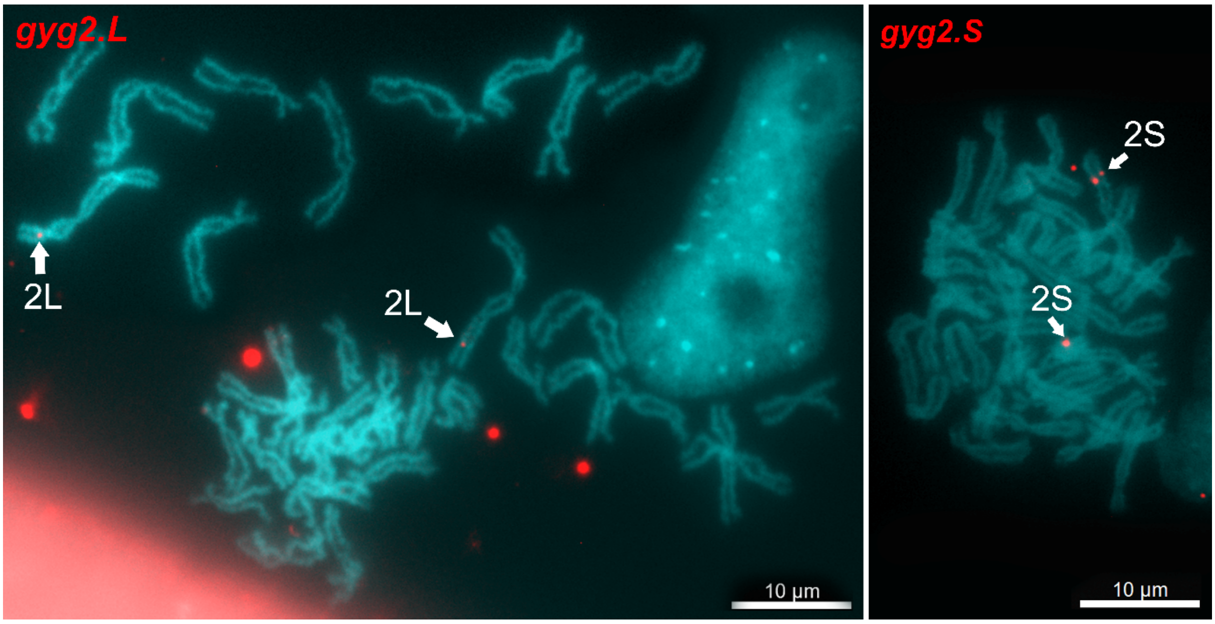


**Figure S6: FISH-TSA with positive *cept1* signals on the *X. borealis* metaphase spread.** The *cept1.L* and *S* genes (red) were localized on the p arm of XBO 2L and 2S, respectively. The *cept1.S* probe hybridized to a single Chr2S gametologue as shown on the metaphase spread on the right side of the panel. Chromosomes were counterstained with DAPI (blue-green). Scale bars represent 10 µm.

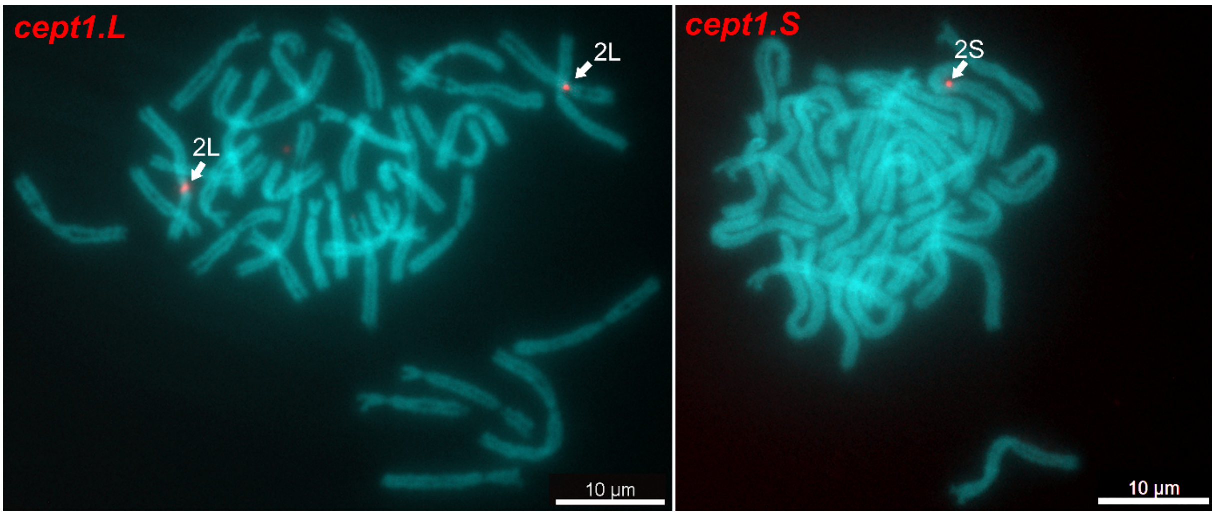

**Figure S7: FISH-TSA with positive *sf3b1* signals on the *X. borealis* metaphase spreads.** The *sf3b1.L* and *S* genes (red) were localized on the q arm of XBO Chr9_10L and 9_10S, respectively. Chromosomes were counterstained with DAPI (blue-green). Scale bars represent 10 µm.

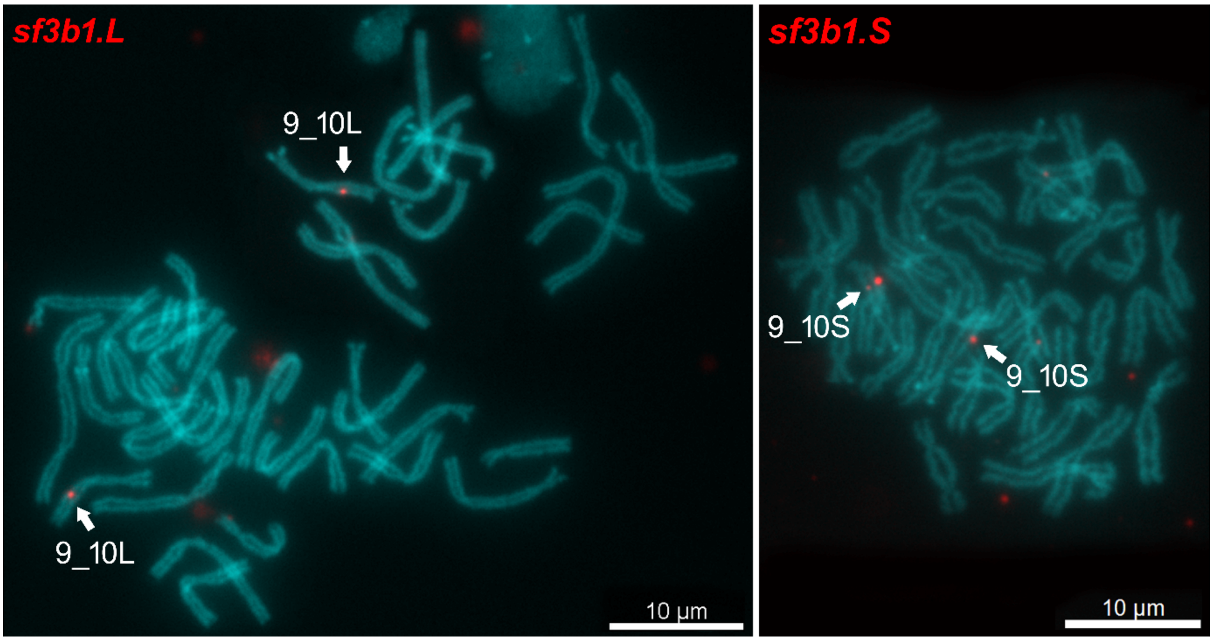


**Figure S8: FISH-TSA with positive *ndufs1* signals on the *X. borealis* metaphase spreads.** The *ndufs1.L* and *S* genes (red) were localized on the q arm of XBO Chr9_10L and 9_10S, respectively. Chromosomes were counterstained with DAPI (blue-green). Scale bars represent 10 µm.

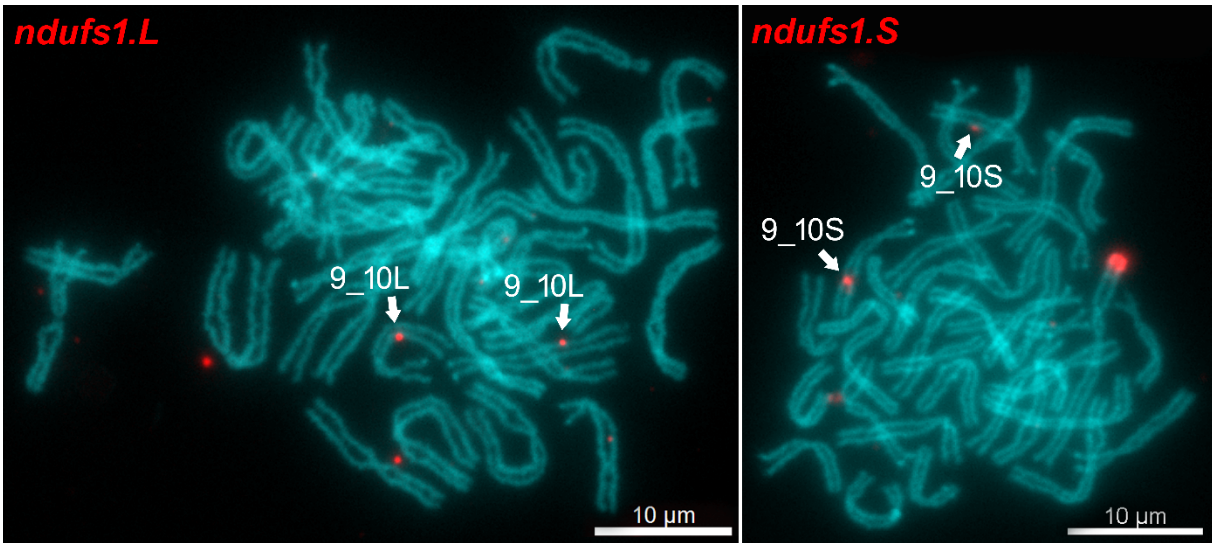

**Figure S9: FISH-TSA with positive *fn1* signals on the *X. borealis* metaphase spreads.** The *fn1.L* and *S* genes (red) were localized on the q arm of XBO Chr9_10L and 9_10S, respectively. Chromosomes were counterstained with DAPI (blue-green). Scale bars represent 10 µm.

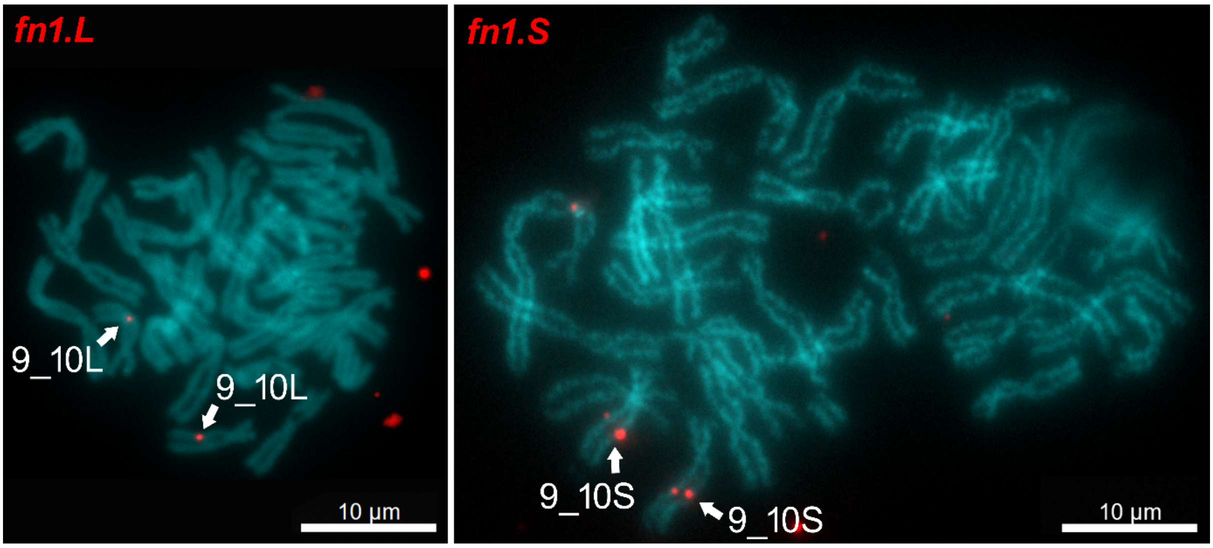


**Figure S10: Whole-genome synteny analysis.** (a) The *X. tropicalis* (Xt) genome containing 10 haploid chromosomes (middle line) aligned with the *X. borealis* L-subgenome (xb L; top) and S-subgenome (xb S; bottom). (b) Similar alignment of the *X. tropicalis* (Xt) genome (middle) with the *X. laevis* L-subgenome (xl L; top) and S-subgenome (xl S; bottom). In both species, the *X. tropicalis* chromosomes serve as the reference for subgenome comparisons.

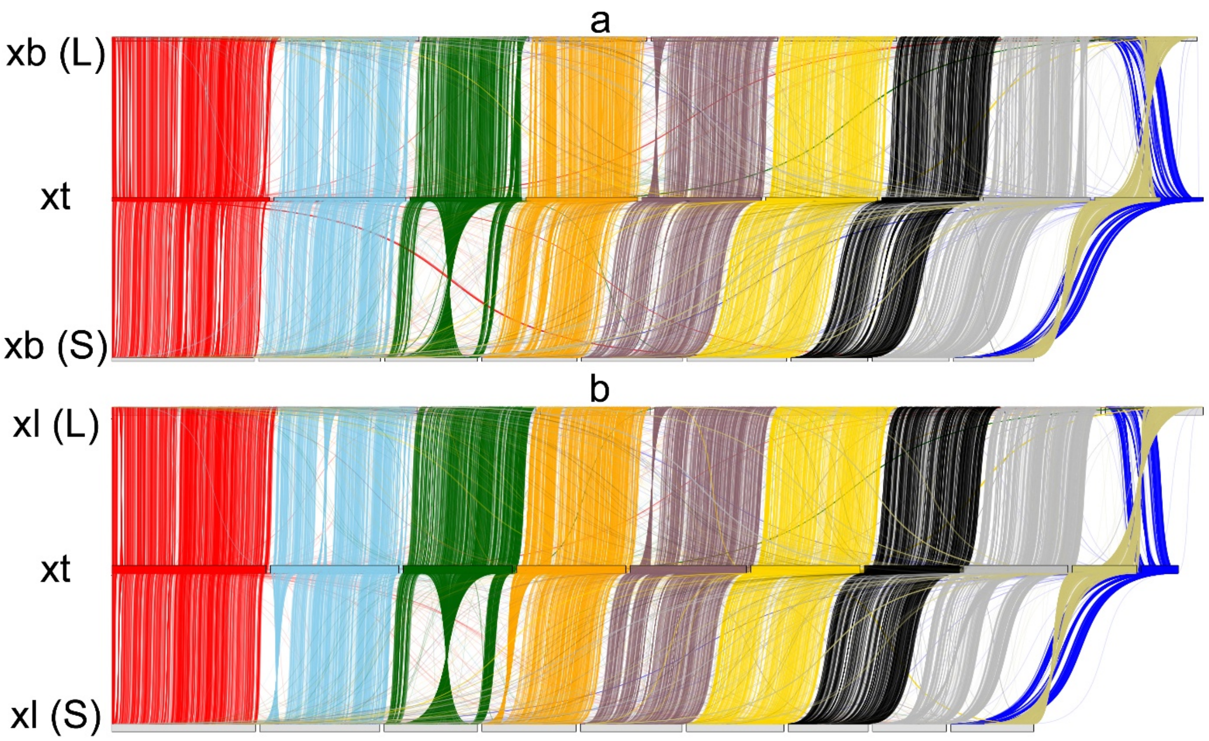
